# Machine learning-guided channelrhodopsin engineering enables minimally-invasive optogenetics

**DOI:** 10.1101/565606

**Authors:** Claire N. Bedbrook, Kevin K. Yang, J. Elliott Robinson, Viviana Gradinaru, Frances H. Arnold

## Abstract

We have engineered light-gated channelrhodopsins (ChRs) whose current strength and light sensitivity enable minimally-invasive neuronal circuit interrogation. Current ChR tools applied to the mammalian brain require intracranial surgery for transgene delivery and implantation of invasive fiber-optic cables to produce light-dependent activation of a small volume of brain tissue [~1 mm^3^]. To enable optogenetics for large brain volumes and without the need for invasive implants, our ChR engineering approach leverages the significant literature of ChR variants to train statistical models for the design of new, high-performance ChRs. With Gaussian Process models trained on a limited experimental set of 102 functionally characterized ChR variants, we designed high-photocurrent ChRs with unprecedented light sensitivity; three of these, ChRger1, ChRger2, and ChRger3, enable optogenetic activation of the nervous system via minimally-invasive systemic transgene delivery with rAAV-PHP.eB, which was not possible previously due to low per-cell transgene copy produced by systemic delivery. These engineered ChRs enable light-induced neuronal excitation without invasive intracranial surgery for virus delivery or fiber optic implantation, i.e. they enable minimally-invasive optogenetics.

## Introduction

Channelrhodopsins (ChRs) are light-gated ion channels found in photosynthetic algae. Transgenic expression of ChRs in the brain enables light-dependent neuronal activation^1^. These channels have been widely applied as tools in neuroscience research^2^; however, functional limitations of available ChRs prohibit a number of optogenetic applications. In their algal hosts, ChRs serve as sunlight sensors in phototaxic and photophobic responses^1^. These channels have broad activation spectra in the visible range and require high-intensity light for activation [~1 mW mm^-2^]. ChRs are naturally low-conductance channels requiring approximately 10^5^ − 10^6^ functional ChRs expressed in the plasma-membrane of a neuron to produce sufficient light-dependent depolarization to induce neuronal activation^3^. When applied to the mouse brain, ChRs require ~1 − 15 mW light delivered ~100 μm from the target cell population to reliably activate action potentials^4–6^. This confines light-dependent activation to a small volume of brain tissue [~1 mm^3^]^7^. Enabling optogenetics for large brain volumes without the need to implant invasive optical fibers for light delivery would be highly desirable for neuroscience applications.

Our goal has been to engineer available ChRs to overcome limits in conductance and light sensitivity and extend the reach of optogenetic experiments. Engineering ChRs requires overcoming three major challenges. First, rhodopsins are trans-membrane proteins that are inherently difficult to engineer because the sequence and structural determinants of membrane protein expression and plasma-membrane localization are highly constrained and poorly understood^8,9^. Second, because properties of interest for neuroscience applications are assayed using low-throughput techniques, such as patch-clamp electrophysiology, engineering by directed evolution is not feasible^10^. And third, *in vivo* applications require either retention or optimization of multiple properties in a single protein tool; for example, we must optimize expression and localization in mammalian cells while simultaneously tuning kinetics, photocurrents, and spectral properties^6^.

Diverse ChRs have been published, including variants discovered from nature^11,12^, variants engineered through recombination^9,13^ and mutagenesis^14,15^, as well as variants resulting from rational design^16^. Studies of these coupled with structural information^17^ and molecular dynamic simulations^18^ have established some understanding of the mechanics and sequence features important for specific ChR properties^1,16^. Despite this, it is still not possible to predict the functional properties of new ChR sequences and therefore not trivial to design new ChRs with a desired combination of functional properties.

Our approach has been to leverage the significant literature of ChRs to train statistical models that enable design of new, highly-functional ChRs. These models take as their input the sequence and structural information for a given ChR variant and then predict its functional properties. The models use training data to learn how sequence and structural elements map to ChR functional properties. Once known, that mapping can be used to predict the functional behavior of untested ChR variants and to select variants predicted to have optimal combinations of desired properties.

We trained models in this manner and found that they accurately predict the functional properties of untested ChR sequences. We used these models to engineer 30 ‘designer’ ChR variants with specific combinations of desired properties. A number of variants identified from this work have unprecedented photocurrent strength and light sensitivity. We have characterized these low-light sensitive, high-photocurrent ChRs for applications in the mammalian brain and demonstrate their potential for minimally-invasive activation of populations of neurons in the brain enabled by systemic transgene delivery with the engineered AAV, rAAV-PHP.eB^19^. This work demonstrates how a machine learning-guided approach can enable engineering of proteins that have been challenging to engineer using existing methods.

## Results

### Dataset of ChR sequence variants and corresponding functional properties for machine learning

In previous work, we explored structure-guided recombination^20,21^ of three highly-functional ChR parents [CsChrimsonR (CsChrimR)^11^, C1C2^17^, and CheRiff^22^] by designing two 10-block recombination libraries with a theoretical size of ~120,000 (i.e. 2×3^10^) ChR variants^9^. Measuring expression, localization, and photocurrent properties of a subset of these chimeric ChRs showed that these recombination libraries are a rich source of functionally diverse sequences^9^. That work produced 76 ChR variants with measured photocurrent properties, the largest single source of published ChR functional data. In subsequent work, we generated an additional 26 ChR variants selected from the same recombination libraries^8^, which we have now characterized for functional properties. Together, these 102 ChR variants from the recombination libraries provide the primary dataset used for model training in this work. We supplemented this dataset with data from other published sources including 19 ChR variants from nature, 14 single-mutant ChR variants, and 28 recombination variants from other libraries (**Dataset 1**). As the data produced by other labs were not collected under the same experimental conditions as data collected in our hands, they cannot be used for comparison for absolute ChR properties (i.e. photocurrent strength); however, these data do provide useful binary information on whether a sequence variant is functional or not. Thus, we used published data from other sources when training binary classification models for ChR function.

Our primary interest was modeling and optimization of three ChR photocurrent properties: photocurrent strength, wavelength sensitivity, and off-kinetics (**Figure 1a**). Enhancing ChR photocurrent strength would enable reliable neuronal activation even under low-light conditions. As metrics of photocurrent strength, we use peak and steady-state photocurrent (**Figure 1a**). As a metric for the ChR activation spectrum, we use the normalized current strength induced by exposure to green light (560 nm) (**Figure 1a**). Different off-rates can be useful for specific applications: fast off-kinetics enable high-frequency optical stimulation^23^, slow off-kinetics is correlated with increased light sensitivity^3,14,15^, and very slow off-kinetics can be used for constant depolarization (step-function opsins [SFOs]^14^). We use two parameters to characterize the off-kinetics: the time to reach 50% of the light-activated current and the photocurrent decay rate, τ_off_ (**Figure 1a**). In addition to opsin functional properties, it is also necessary to optimize or maintain plasma-membrane localization, a prerequisite for ChR function^8^.

**Figure 1.**
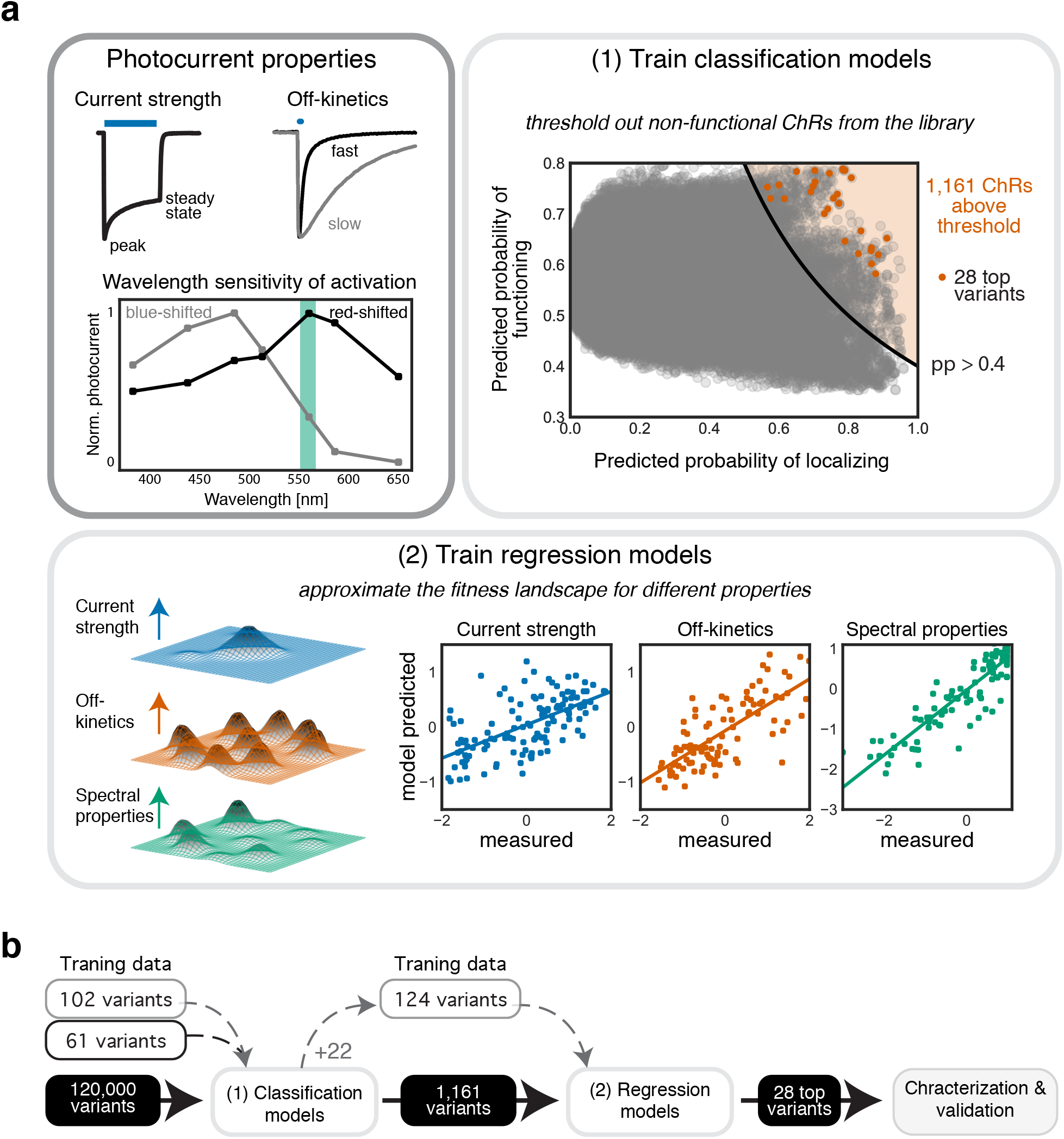
Machine learning-guided optimization of ChR photocurrent strength, off-kinetics, and wavelength sensitivity of activation. (**a**) Upon light exposure, ChRs open and reach a *peak* inward current; with continuous light exposure, ChRs desensitize reaching a lower *steady-state* current. We used both peak and steady-state current as metrics for photocurrent strength. To evaluate ChR off-kinetics we used the current decay rate (τ_off_) after a 1 ms light exposure and also the time to reach 50% of the light-exposed current after light removal. ChRs are maximally activated by one wavelength of light. ‘Blue shifted’ ChRs have a peak activation wavelength between ~450-480 nm, while ‘red shifted’ ChRs have a peak activation wavelength between 520-650 nm. We used the normalized photocurrent with green (560 nm) light as a metric for wavelength sensitivity of activation. For variant selection, we trained classification models to predict whether ChRs would localize correctly to the plasma membrane and function (**1**) and then trained regression models to approximate the fitness landscape for each property of interest for the recombination library (**2**). Sequences predicted to localize and function by the classification models and predicted to have an optimized set of functional properties by the regression models were selected for further characterization, e.g., the 28 top variants. Models were trained with photocurrent properties for each ChR in the training set (plots show 20-fold cross validation on the training set). (**b**) The classification function model was trained with 102 recombination variants (**Dataset 2**) and 61 previously-published ChRs (**Dataset 1**) and the regression models were trained with 124 recombination variants (**Dataset 2**).

As inputs for the machine-learning models, we consider both ChR sequence and structure. ChR sequence information is simply encoded in the amino acid sequence. For structural comparisons, we convert the 3D crystal-structural information into a ‘contact map’ that is convenient for modeling. Two residues are considered to be in contact and potentially important for structural and functional integrity if they have any non-hydrogen atoms within 4.5 Å in the C1C2 crystal structure^17^.

### Training Gaussian process (GP) classification and regression models

Using the ChR sequence/structure and functional data as inputs, we trained Gaussian process (GP) classification and regression models (**Figure 1**). GP models have successfully predicted thermostability, substrate binding affinity, and kinetics for several soluble enzymes^24^, and, more recently, ChR membrane localization^8^. For a detailed description of the GP model architecture and properties used for protein engineering see refs 8, 23. Briefly, these models infer predictive values for new sequences from training examples by assuming that similar inputs (ChR sequence variants) will have similar outputs (photocurrent properties). To quantify the relatedness of inputs (ChR sequence variants), we compared both sequence and structure. We defined the sequence and structural similarity between two chimeras by aligning them and counting the number of positions and contacts at which they are identical^24^.

We trained a binary classification model to predict if a ChR sequence will be functional using all 102 training sequences from the recombination library as well as data from 61 sequence variants published by others (**Dataset 1**). A ChR sequence was considered to be functional if its photocurrents were >0.1 nA upon light exposure, a threshold we set as an approximate lower bound for current necessary for neuronal activation. We then used this trained classification model to predict whether uncharacterized ChR sequence variants were functional (**Figure 1a**). To verify that the classification model is capable of accurate predictions, we performed 20-fold cross validation on the training data set and measured an area under the receiver operator curve (AUC) of 0.78, indicating good predictive power (**Table 1**).

**Table 1.** Evaluation of prediction accuracy for different ChR property models. Calculated AUC or Pearson correlation after 20-fold cross validation on training set data for classification and regression models. The test set for both the classification and regression models was the 28 ChR sequences predicted to have useful combinations of diverse properties. Accuracy of model predictions on the test set is evaluated by AUC (for classification model) or Pearson correlation (for the regression models). The Matérn kernel is with 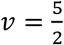.

| Model type | ChR property | Kernel | Cross validation | Test set |
| --- | --- | --- | --- | --- |
| GP classification | function | Matérn | AUC = 0.78 | AUC = 1.0 |
| GP regression | current strength | Matérn | R = 0.77 | R = 0.92 |
| GP regression | off-kinetics | Matérn | R = 0.78 | R = 0.97 |
| GP regression | wavelength sensitivity | Matérn | R = 0.89 | R = 0.96 |

Next, we trained three regression models, one for each of the ChR photocurrent properties of interest: photocurrent strength, wavelength sensitivity of photocurrents, and off-kinetics (**Figure 1a**). For these, we exclusively used data collected from our ChR recombination libraries (**Dataset 2**). Once trained, these models were used to predict photocurrent strength, wavelength sensitivity of photocurrents, and off-kinetics of new, untested ChRs sequence variants. Again, to test whether these models make accurate predictions, we performed 20-fold cross validation on the training dataset and observed high correlation between predicted and measured properties as indicated by Pearson correlations between 0.77 − 0.89 for all models (**Table 1**).

### Selection of designer ChRs using trained models

A ‘designer’ ChR is a ChR predicted by the models to have a useful combination of properties. We used a tiered approach (**Figure 1b**) to select designer ChRs. The first step was to eliminate all ChR sequences predicted to not localize to the plasma membrane or predicted to be nonfunctional. To do this, we used the ChR function classification model along with the previously published ChR localization classification model^8^ to predict the probability of localization and function for each ChR sequence in the 120,000-variant recombination library. Not surprisingly, most ChR variants were predicted to not localize and not function. To focus on ChR variants predicted to localize and function, we set a threshold for the product of the predicted probabilities of localization and function; any ChR sequence above that threshold would be considered for the next tier of the process. We selected a conservative threshold of 0.4.

The model training data made clear that the higher the mutation level (mutation distance from one of the three parent proteins), the less likely it was that a sequence would be functional; however, we expect that more diverse sequences would also have the more diverse functional properties. We wanted to explore diverse sequences predicted to function by the classification models. We selected 22 ChR variants that passed the 0.4 threshold and were diverse multi-block-swap sequences (i.e. containing on average 70 mutations from the closest parent). After these 22 sequences were synthesized, cloned in the expression vector, and expressed in HEK cells, their photocurrent properties were measured with patch-clamp electrophysiology. 59% of the tested sequences were functional (**Figure 2a**), compared to 38% of the multi-block swap sequences not selected by the model and having the same average mutation level. This validates the classification model’s ability to make useful predictions about novel functional sequences, even for sequences that are very distant from those previously tested. We then updated the models by including data from these 22 sequences for future rounds of predictions.

**Figure 2.**
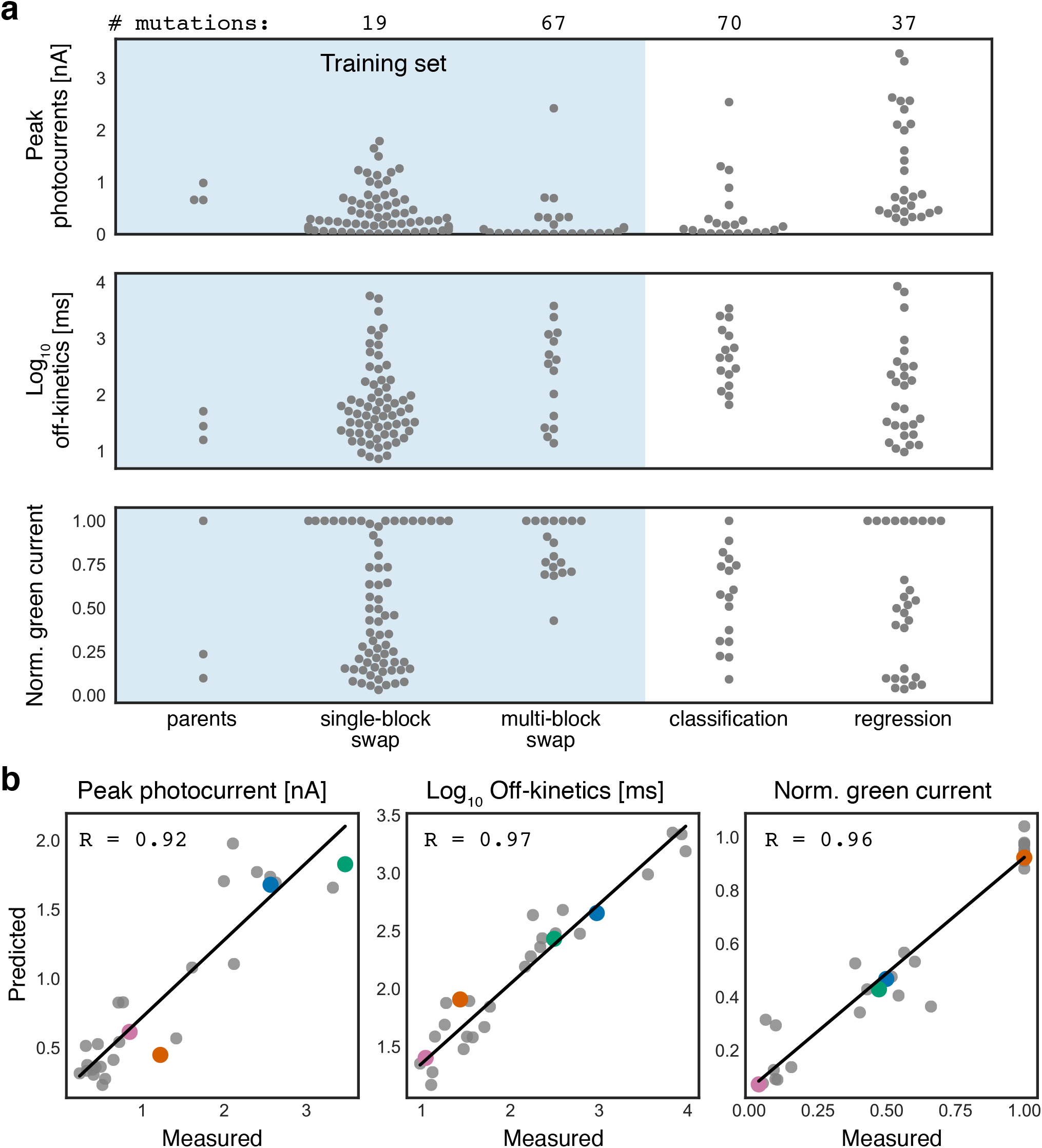
Training machine-learning models to predict ChR properties of interest based on sequence and structure enables design of ChR variants with collections of desirable properties. (**a**) Measurements of training set ChR and model-predicted ChR, peak photocurrent, off-kinetics, and normalized green current. Each gray-colored point is a ChR variant. Training set data are shaded in blue. Mean number of mutations for each set is above the plots. (**b**) Model predictions vs measured property for peak photocurrent, off-kinetics, and normalized green current of the 28 designer ChRs shows strong correlation. Specific ChR variants are highlighted to show predicted and measured properties for all three models: blue, ChR_12_10, green, ChR_11_10, orange, ChR_28_10, pink, ChR_5_10.

Of the 120,000-variant recombination library, 1,161 chimeric sequence variants passed the conservative 0.4 predicted localization and function threshold (**Figure 1**). For the second tier of the selection process, we used the three regression models trained on all functional variants collected up to this point to predict the photocurrent strength, wavelength sensitivity of photocurrents, and off-kinetics for each of these 1,161 ChR sequence variants (**Dataset 3**). From these predictions, we selected those ChRs predicted to have the highest photocurrent strength, most red-shifted or blue-shifted activation wavelengths, and those with a range of off-kinetics from very fast to very slow. We selected 28 designer ChRs with different combinations of properties that were all predicted to be highly functional (photocurrents > 0.2 nA) and capable of good membrane localization (**Supplemental Figure 1-2**).

Genes encoding the 28 selected designer ChR variants were synthesized and cloned into expression vectors, expressed in HEK cells, and characterized for their photocurrent properties with patch-clamp electrophysiology. All 28 selected designer ChRs were functional: 100% of chimeras selected using the updated classification model above the 0.4 threshold both localize and function. For each of the designer ChR variants, the three measured photocurrent properties correlated very well with the model predictions (R>0.9 for all models) (**Figure 2b**, **Table 1**). This outstanding performance on a novel set of sequences demonstrates the power of this data-driven predictive method for engineering designer ChRs. As a negative control, we selected two ChR variant sequences from the recombination library that the model predicted would be nonfunctional (ChR_29_10 and ChR_30_10). These sequences resulted from a single-block swap from two of the most highly functional ChR recombination variants tested. As predicted, these sequences were non-functional (**Figure 3b**), which shows that ChR functionality can be attenuated by incorporating even minimal diversity at certain positions.

**Figure 3.**
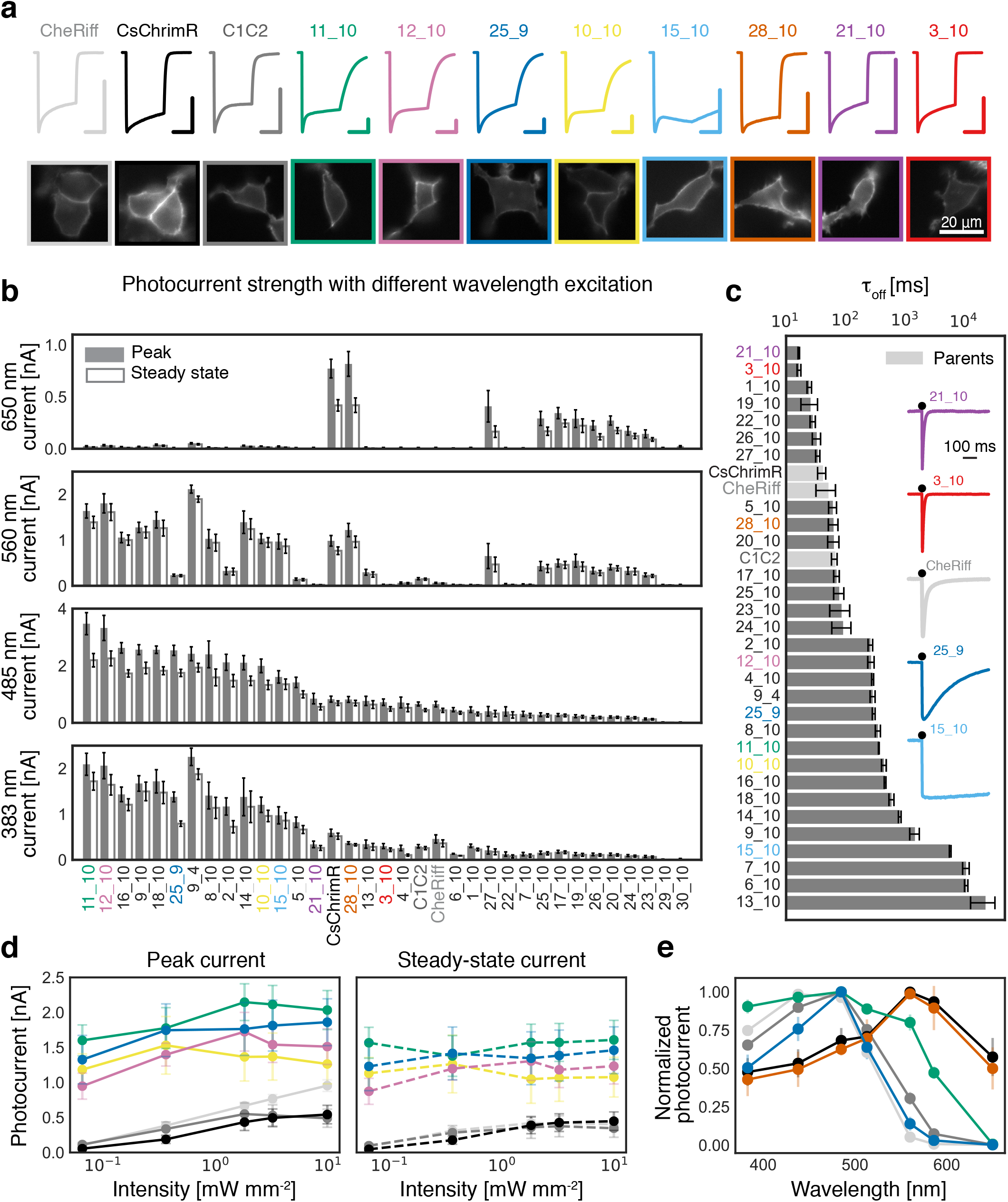
The model-predicted ChRs exhibit a large range of functional properties often far exceeding the parents. (**a**) Current trace after 0.5 s light exposure for select designer ChR variants with corresponding expression and localization in HEK cells. Vertical colored scale bar for each ChR current trace represents 0.5 nA, and horizontal scale bar represents 250 ms. Different color traces are labeled with each variant’s name. The variant color presented in (**a**) is kept constant for all other panels. (**b**) Designer ChR measured peak and steady-state photocurrent with different wavelengths of light in HEK cells (*n* = 4–8 cells, see **Dataset 2**). 383 nm light at 1.5 mW mm^-2^, 485 nm light at 2.3 mW mm^-2^, 560 nm light at 2.8 mW mm^-2^, and 650 nm light at 2.2 mW mm^-2^. (**c**) Designer ChR off-kinetics decay rate (τ_of_f) following a 1 ms exposure to 485 nm light at 2.3 mW mm^-2^ (*n* = 4–8 cells, see **Dataset 2**). Parent ChRs are highlighted in light gray. Inset shows current traces with 1 ms light exposure for select ChRs compared with CheRiff. (**d**) Selected ChR variants’ peak and steady-state photocurrent strength with varying light irradiances compared with parental ChRs (CheRiff, *n* = 5; CsChrimR, *n* = 5; C1C2, *n* = 4; 28_10, *n* = 5; 11_10, *n* = 5; 25_9, *n* = 5). (**e**) Wavelength sensitivity of activation for select ChRs compared with parental ChRs (CheRiff, *n* = 6; CsChrimR, *n* = 5; C1C2, *n* = 4; 11_10, *n* = 6; 12_10, *n* = 7; 25_9, *n* = 5; 10_10, *n* = 4). Top variants, ChR_9_4, ChR_25_9, and ChR_11_10 are named ChRger1, ChRger2, and ChRger3 in subsequent figures. Plotted data are mean ± SEM.

### Sequence and structural determinants of ChR functional properties

We used L1-regularized linear regression models to identify a limited set of residues and structural contacts that strongly influence ChR photocurrent strength, spectral properties, and off-kinetics. We can assess the relative importance of these sequence and structural features by weighting their contributions using L2-regularized linear regression and have included important features and their weights in **Dataset 4** and **Supplemental Figure 3-4**. For each functional property, we identified a set of important residues and contacts. Residues and contacts most important for tuning spectral properties are generally proximal to the retinal-binding pocket, with some exceptions (**Supplemental Figure 4**). Residues important for photocurrent strength reside between transmembrane (TM) helix 1 and 7 (**Supplemental Figure 3**). The C1C2 crystal structure shows TM helices 1, 2 and 7 form a cavity which allows water influx for the cation-translocation pathway. Interestingly, residues important for photocurrent strength also appear to be important for kinetic properties (**Supplemental Figure 3**), consistent with previous findings that light sensitivity is inversely proportional to off-kinetic speed^3,14,15^.

### Machine-guided search identifies ChRs with a range of useful functional properties

We assessed photocurrent amplitude, wavelength sensitivity, and off-kinetics of the designer ChRs and the three parental ChRs (CsChrimR^11^, CheRiff^22^, and C1C2^17^) (**Figure 3**). In addition to the 28 regression model-predicted ChRs, we also assessed the top performing ChRs from the classification models’ predictions [ChR_9_4 (predicted from the classification localization model) and ChR_25_9 (classification function model)], for a total of 30 highly-functional model-predicted ChRs as well as the two negative control ChRs (ChR_29_10, ChR_30_10). Of the 30 model-predicted ChRs, we found 12 variants with ≥2-times higher blue-light activated photocurrents than the top-performing parent (CsChrimR) (**Figure 3b**). Three variants exhibit ≥1.7-times higher green-light activated photocurrents than CsChrimR (**Figure 3b**). Eight variants have larger red-light activated photocurrents when compared with the blue-light activated parents (CheRiff and C1C2), though none out-perform CsChrimR (**Figure 3b**). Both ChR variants predicted to be non-functional by the models produce <0.03 nA currents.

Characterization of the 30 designer ChRs revealed that their off-kinetics span three orders of magnitude (τ_off_ =10 ms − >10 s) (**Figure 3c**). This range is quite remarkable given that all designer ChRs are built from sequence blocks of three parents that have very similar off-kinetics (τ_off_ = 30 − 50 ms). We found that 5 designer ChRs have faster off-kinetics than the fastest parent, while 16 have >5-times slower off-kinetics (**Figure 3c**). The two fastest variants, ChR_3_10 and ChR_21_10 exhibit τ_off_ = 13 ± 0.9 ms and 12 ± 0.4 ms, respectively (mean ± SEM). Four ChRs have particularly slow off-kinetics with τoff >1 s, including ChR_15_10, ChR_6_10, and ChR_13_10 (τ_off_ = 4.3 ± 0.1 s, 8.0 ± 0.5 s, and 17 ± 7 s, respectively). Two ChRs with very strong photocurrents, ChR_25_9 and ChR_11_10, exhibit τ_off_ = 220 ± 10 ms and 330 ± 30 ms, respectively. Short 1 ms-exposures to blue light elicits distinct profiles from selected ChRs: ChR_21_10 turns off rapidly, ChR_25_9 and ChR_11_10 turn off more slowly, and ChR_15_10 exhibits little decrease in photocurrent 0.5 s after the light was turned off (**Figure 3c**).

Three designer ChRs exhibit interesting spectral properties. ChR_28_10’s red-shifted spectrum matches that of CsChrimR, demonstrating that incorporating sequence elements from blue-shifted ChRs into CsChrimR can still generate a red-shifted activation spectrum (**Figure 3e**). Two of the designer ChRs exhibit novel spectral properties: ChR_11_10 has a broad activation spectrum relative to the parental spectra, with similar steady-state current strength from 400 − 560 nm light and even maintain strong currents (0.7 ± 0.1 nA) when activated with 586 nm light (**Figure 3e**). ChR_25_9, on the other hand, exhibits a narrow activation spectrum relative to the parental spectra, with a peak at 485 nm light (**Figure 3e**).

We assessed the light sensitivity of the designer ChRs with enhanced photocurrents by measuring photocurrent strength at various irradiances (**Figure 3d**). We refer to these high-photocurrent ChRs as ‘high-performance’ ChRs. Compared with CsChrimR, CheRiff, and C1C2, all high-performance ChRs have ≥9-times larger currents at the lowest intensity of light tested (10^-1^ mW mm^-2^) as well as larger currents at all intensities of light tested. The high-performance ChRs also demonstrate minimal decrease in photocurrent magnitude over the range of intensities tested (10^-1^ − 10^1^ mW mm^-2^), suggesting that photocurrents were saturated at these intensities and would only attenuate at much lower light intensities (**Figure 3d**). The high-performance ChRs are expressed at levels similar to the CsChrimR parent (the highest expressing parent) indicating that the improved photocurrent strength of these ChRs is not solely due to improved expression (**Supplemental Figure 5-6**).

We also compared high-performance designer ChRs with ChR2(H134R)^6,25^, an enhanced photocurrent single mutant of ChR2 commonly used for *in vivo* optogenetics, and CoChR (from *Chloromonas oogama)^11^*, which was reported to be one of the highest conducting ChRs activated by blue light. Three of the top high-performance ChRs (ChR_9_4, ChR_25_9, and ChR_11_10) show significantly larger peak and steady-state currents compared with ChR2 and significantly larger steady-state currents when compared with CoChR when exposed to 2 mW mm^-2^ 485 nm light (**Supplemental Figure 7d,f**). Although CoChR produced peak currents of similar magnitude to the high-performance ChRs, CoChR decays to a much lower steady-state level (**Supplemental Figure 7d,f**). At lower light intensities (6.5×10^-2^ mW mm^-2^), the high-performance ChRs produce significantly larger photocurrents than both ChR2(H134R) and CoChR (**Supplemental Figure 7e,g**). These high-performance opsins have the potential for optogenetic activation with very low light levels. The increased low-light sensitivity of these high-performance ChRs is likely due in part to their relatively slow off kinetics leading to the increased accumulation of the open state under low-light conditions^14^.

### Validation of designer ChRs for neuroscience applications

For further validation in neurons we selected three of the top high-conductance ChRs, ChR_9_4, ChR_25_9, and ChR_11_10, and renamed them ChRger1, ChRger2, and ChRger3, respectively, for **<u>ch</u>**annel**<u>r</u>**hodopsin **<u>G</u>**aussian process-**<u>e</u>**ngineered **<u>r</u>**ecombinant opsin (**Supplemental Figure 8**). For validation in cultured neurons and acute brain slices, ChRger1-3 and ChR2(H134R) were cloned into AAV vectors with either a hSyn or CaMKIIa promoter, Golgi export trafficking signal (TS) sequence^5^, and enhanced yellow fluorescent protein (eYFP) marker and packaged in the engineered rAAV-PHP.eB capsid^19^ (**Figure 4a** and **Supplemental Table 1**). When expressed in cultured neurons under the hSyn promoter, the ChRgers display robust membrane localization and expression throughout the neuron soma and neurites (**Figure 4a**). We assessed neuronal spike fidelity with varying irradiance using ChR2(H134R) for comparison and observed a 10 − 100-fold decrease in the light intensity required to induce reliable spiking by 1 and 5 ms 485 nm light pulses (**Figure 4b,c**). These results demonstrate that the designer ChRgers require 1 − 2 orders of magnitude lower light intensity than ChR2(H134R) for neuronal activation.

**Figure 4.**
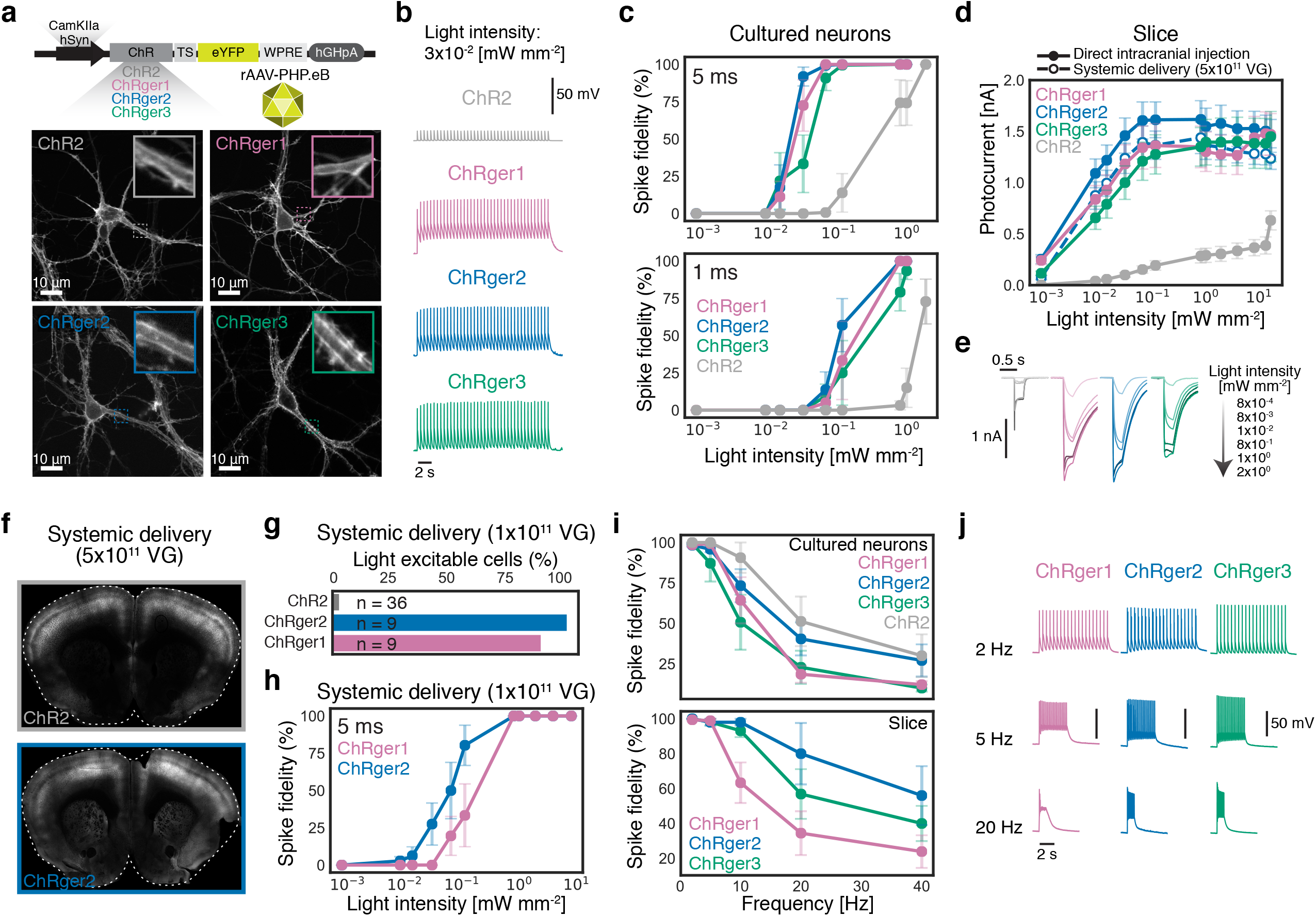
ChRger variants in cultured neurons and in acute brain slices outperform the commonly used ChR2(H134R). (**a**) ChRgers and the ChR2(H134R) control were cloned into an AAV vector with either the hSyn or CamKIIa promoter, a trafficking signal (TS), eYFP, and WPRE and then packaged into rAAV-PHP.eB for expression in culture and *in vivo*. Cultured neurons expressing ChRgers and ChR2(H134R) under the hSyn promoter. (**b**) Voltage traces of ChRgers and ChR2(H134R) at 2 Hz with 5 ms pulsed low-intensity blue light stimulation (3×10^-2^ mW mm^-2^) shows robust neuronal firing for ChRgers while ChR2(H134R) exhibits only sub-threshold light-induced depolarization. (**c**) Spike fidelity with varying intensity light of ChRgers and ChR2(H134R) for 5 ms and 1 ms light-pulse width at 2 Hz stimulation (ChRger1, *n* = 6; ChRger2, *n* = 4; ChRger3, *n* = 6; ChR2, *n* = 7). (**d**) ChRgers and ChR2(H134R) photocurrent strength with varying light irradiances in acute brain slice after direct injection of rAAV-PHP.eB packaged hSyn-ChR constructs into the PFC (ChRger1, *n* = 11; ChRger2, *n* = 11; ChRger3, *n* = 11; ChR2, *n* = 9) or after systemic delivery of CamKIIa-ChRger2 (ChRger2, *n* = 6; 5×10^11^ vg/animal). (**e**) Current traces of ChRgers and ChR2(H134R) with a 300 ms light pulse at varying light irradiances in acute brain slice after direct injection. (**f**) Systemic delivery of rAAV-PHP.eB packaged hSyn-ChRger2 or hSyn-ChR2(H134R) resulted in broad expression throughout the cortex (5×10^11^ vg/animal). (**g**) The fraction of light excitable neurons in the PFC after systemic delivery of rAAV-PHP.eB packaged hSyn-ChRgers or hSyn-ChR2(H134R) measured by cell-attached recording in acute slice targeting only neurons expressing the eYFP marker (1×10^11^ vg/animal). (**h**) Spike fidelity with varying intensity light of hSyn-ChRgers after systemic delivery (1×10^11^ vg/animal) (ChRger1, *n* = 5; ChRger2, *n* = 8). (**i**) Spike fidelity with varying stimulation frequency of hSyn-ChRgers in cultured neurons (top) with 2 ms light-pulse width (ChRger1, *n* = 9; ChRger2, *n* = 12; ChRger3, *n* = 7; ChR2, *n* = 8), or in acute brain slice after systemic delivery of CamKIIa-ChRgers (bottom; 1×10^11^ vg/animal) with 0.5 ms light-pulse width (ChRger1, *n* = 9; ChRger2, *n* = 5; ChRger3, *n* = 8). Spike fidelity in culture and in slice was done with 1 mW mm^-2^ intensity light. (**j**) Voltage traces with blue light–driven spiking at the indicated frequencies with 1 mW mm^-2^. vg, viral genomes. Plotted data are mean ± SEM.

Next, we performed direct intracranial injections into the mouse prefrontal cortex (PFC) of rAAV-PHP.eB packaging either ChRger1, ChRger2, ChRger3, or ChR2(H134R) under the hSyn promoter. After 3 − 5 weeks of expression, we measured light sensitivity in ChR-expressing neurons in acute brain slices. Consistent with the pervious experiments, we observed a large increase in the light sensitivity for the ChRgers compared with ChR2(H134R) (**Figure 4d,e**). The ChRgers exhibit >200 pA photocurrent at the lowest irradiance tested, 10^-3^ mW mm^-2^, while at the equivalent irradiance ChR2(H134R) exhibits undetectable photocurrents (**Figure 4d,e**). The ChRgers reach >1 nA photocurrents with ~10^-2^ mW mm^-2^ light, a four-fold improvement over ChR2(H134R)’s irradiance-matched photocurrents (**Figure 4d**). Our characterization of ChR2(H134R)’s light sensitivity and photocurrent strength is consistent with previously published results from other labs^6,22^.

### Designer ChRs and systemic AAVs enable minimally-invasive optogenetic excitation

We investigated whether these light-sensitive, high-photocurrent ChRs could provide optogenetic activation coupled with minimally-invasive gene delivery. Previous reports of ‘non-invasive optogenetics’ relied on invasive intracranial virus delivery, which results in many copies of virus per cell and thus very high expression levels of the injected construct^26^. Recently, we described the novel AAV capsid rAAV-PHP.eB^19^ that produces broad transduction throughout the central nervous system with a single minimally-invasive intravenous injection in the adult mouse^27,28^. Systemic delivery of rAAV-PHP.eB vectors results in brain-wide transgene delivery with expression throughout large brain volumes without the need for invasive intracranial injections^19,27,28^. The use of rAAV-PHP.eB for optogenetic applications has been limited, however, by the lower multiplicity of infection with systemically delivered viral vectors than with direct injection. This results in insufficient opsin expression and light-evoked currents to evoke neuronal firing with commonly-used channels (e.g. ChR2).

We hypothesized that the ChRgers could overcome this limitation and allow large-volume optogenetic excitation following systemic transgene delivery. We systemically delivered rAAV-PHP.eB packaging either ChRger1-TS-eYFP, ChRger2-TS-eYFP, or ChR2(H134R)-TS-eYFP under the hSyn promoter and observed broad expression throughout the brain with expression strongest in the cortex (**Figure 4f**). We then measured the fraction of opsin-expressing cells with sufficient opsin-mediated currents for light-induced firing (**Figure 4g**). Only 1/36 neurons expressing ChR2(H134R) produced light-induced firing, while 8/9 neurons expressing ChRger1 produced light-induced activity and 9/9 neurons expressing ChRger2 produced light-induced activity. We also observed high spike fidelity with low light levels in ChRger1 and ChRger2, consistent with observations in neuronal cultures (**Figure 4h**). These results demonstrate the need for light-sensitive and high-photocurrent opsins for applications where systemic delivery is desired.

We also systemically delivered rAAV-PHP.eB packaging ChRger1-3 under the CaMKIIa promoter. With systemic delivery of ChRger2, we observed photocurrent strength similar to results observed after direct injection into the PFC (**Figure 4d**). When expressed in pyramidal neurons in the cortex, ChRger2 and ChRger3 enabled robust optically-induced firing at rates between 2 − 10 Hz, although spike fidelity was reduced at higher frequency stimulation (**Figure 4i,j**). ChRger2 performed best with higher frequency stimulation while ChRger1 performed worst. The ChRgers also produced robust light-induced spiking with short pulse-width stimulation (e.g., 0.5 ms pulse width; Figure 4i).

We next evaluated the optogenetic efficiency of ChRger2 after systemic delivery using a well-established behavioral paradigm: optogenetic intracranial self-stimulation (oICSS) of dopaminergic neurons of the ventral tegmental area (VTA)^29^. We used systemic delivery of rAAV-PHP.eB packaging a double-floxed inverted open reading frame (DIO) containing either ChRger2-TS-eYFP or ChR2(H134R)-TS-eYFP into *Dat*-Cre mice (**Figure 5a** and **Supplemental Table 1**). Three weeks after systemic viral delivery and stereotaxic implantation of fiber-optic cannulas above the VTA, mice were placed in an operant box and were conditioned to trigger a burst of 447 nm laser stimulation via nose poke. Animals expressing ChRger2 displayed robust optogenetic self-stimulation in a frequency-dependent and laser power-dependent manner. Higher frequencies (up to 20 Hz) and higher light power (up to 10 mW) promoted greater maximum operant response rates (**Figure 5a**). Conversely, laser stimulation failed to reinforce operant responding in ChR2(H134R)-expressing animals (**Figure 5a**); these results were consistent with results in acute slice where the light-induced currents of ChR2(H134R) are too weak at the low copy number produced by systemic delivery for robust neuronal activation.

**Figure 5.**
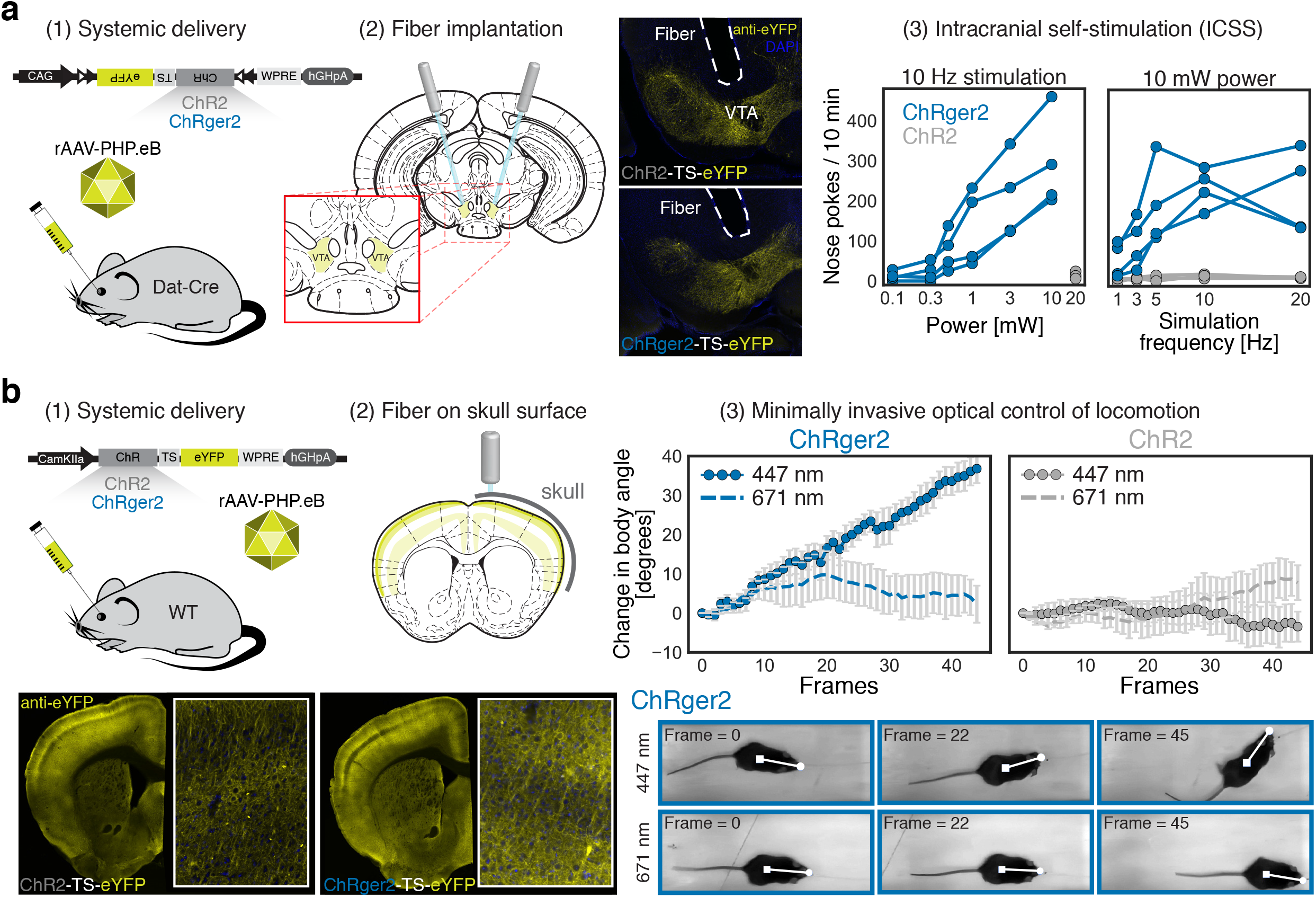
Validation of high-performance ChRger2 for minimally-invasive optogenetic behavioral modulation. (**a**) Minimally-invasive, systemic delivery of rAAV-PHP.eB packaged CAG-DIO ChRger2-TS-eYFP or ChR2(H134R)-TS-eYFP (3×10^11^ vg/mouse) into *Dat*-Cre animals coupled with fiber optic implantation above the VTA enabled blue light-induced intracranial self-stimulation (ten 5 ms laser pulses) exclusively with ChRger2 and not ChR2(H134R) with varying light power and varying stimulation frequencies. ChRger2, *n* = 4; ChR2(H134R), *n* = 4. Images show fiber placement and opsin expression for ChR2(H134R) (top) and ChRger2 (bottom). (**b**) Minimally-invasive, systemic delivery of rAAV-PHP.eB packaged CaMKIIa ChRger2-TS-eYFP or ChR2(H134R)-TS-eYFP (5×10^11^ vg/mouse) into wild type (WT) animals coupled with surgically secured 2 mm long, 400 μm fiber-optic cannula guide to the surface of the skull above the right M2 that had been thinned to create a level surface for the fiber-skull interface. Three weeks later, mice were trained to walk on a linear-track treadmill at fixed velocity. Coronal slices show expression throughout cortex with higher magnification image of M2 (inset) for ChR2(H134R) (left) and ChRger2 (right). Unilateral blue light stimulation of M2 induced turning behavior exclusively with ChRger2 and not ChR2(H134R) (10 Hz stimulation with 5 ms 447 nm light pulses at 20 mW). ChRger2, *n* = 5; ChR2(H134R), *n* = 5. No turning behavior was observed in any animal with 10 Hz stimulation with 5 ms 671 nm light pulses (20 mW). Plotted data are mean ± SEM. vg, viral genomes.

In order to determine if ChRger2 would enable both minimally-invasive transgene delivery and minimally-invasive optical excitation, we assayed directional control of locomotion in freely moving animals by optogenetic stimulation of the right secondary motor cortex (M2), a well-established behavioral paradigm previously used to validate optogenetic tools^30^. In this assay, unilateral stimulation of M2 disrupts motor function in the contralateral lower extremities, causing mice to turn away from the stimulation side. We systemically administered rAAV-PHP.eB packaging either ChRger2-TS-eYFP or ChR2(H134R)-TS-eYFP under a CaMKIIa promoter for transgene expression in excitatory pyramidal neurons in the cortex (**Figure 5b**, and **Supplemental Table 1**). We observed broad expression throughout the cortex for both ChRger2 and ChR2(H134R) injected animals (**Supplemental Figure 9**). We secured a fiber-optic cannula guide to the surface of the thinned skull above M2 without puncturing the dura and therefore leaving the brain intact (**Figure 5b**), which we consider to be minimally invasive. Despite the presence of the highly optically scattering calavarial bone, stimulation with 20 mW 447 nm light induced left-turning behavior in animals expressing ChRger2 but not in animals expressing ChR2(H134R) (**Figure 5b** and **Supplemental Video 1-2**). Left-turning behavior terminated upon conclusion of optical stimulation (**Supplemental Video 1**). Behavioral effects were seen at powers as low as 10 mW, but the most consistent turning phenotypes were seen with 20 mW laser power. In order to ensure that turning behavior was not due to unexpected visual stimuli or heating caused by the stimulation laser, we repeated treadmill experiments using 671 nm light, which is outside the excitation spectrum of both opsins. 20 mW 671 nm light failed to induce turning in both ChRger2 and ChR2(H124R). Overall, these experiments demonstrate that ChRger2 is compatible with minimally-invasive systemic gene delivery and can enable minimally-invasive optogenetic excitation.

## Discussion

We have outlined and demonstrated a data-driven approach to engineering ChR properties that enables efficient discovery of highly functional ChR variants based on data from relatively few variants. In this approach we approximate the ChR fitness landscape and use it to efficiently search sequence space and select top-performing variants for a given property^10,24^. By first eliminating the vast majority of non-functional sequences, we can focus on local peaks scattered throughout the landscape. Then, using regression models, we predict which sequences lie on the fitness peaks.

Designing useful ChRs for *in vivo* applications requires simultaneous optimization of multiple properties; machine learning provides a platform for such optimization and allows us to identify designer variants with combinations of properties that follow engineering specifications. Using a limited sequence space of ~120,000 chimeric ChRs, we were able to generate variants with large variations in off-kinetics (10 ms to >10 s) and photocurrents that far exceed any of the parental or other commonly used ChRs. We also use the machine-learning models to identify the residues and contacts most important for ChR function. Application of this machine-learning pipeline (limited data collection from diverse sequences, model training and validation, and prediction and testing of new sequences) is likely to generate other new and improved protein-based neuroscience tools, e.g., anion-conducting ChRs^12^, calcium sensors, voltage sensors^31^, and AAVs^27^.

We have designed high-performance ChRs (ChRger1, ChRger2, and ChRger3) with unprecedented light sensitivity and have validated ChRger2’s application for *in vivo* optogenetics. The high-photocurrent properties of these ChRs have overcome the limitation of low per-cell copy number after systemic delivery. ChRger2 enabled neuronal excitation with high temporal precision without invasive intracranial surgery for virus delivery or fiber optic implantation for superficial brain areas, extending what is currently possible for optogenetics experiments. Coupling ChRgers with recently reported upconversion nanoparticles may allow for non-invasive optogenetics in deep brain areas with systemic transgene delivery and tissue-penetrating near-infrared (NIR) light for neuronal excitation^26^.

### Online methods

#### Construct design and characterization

The design, construction, and characterization of the recombination library of chimeras is described in detail in Bedbrook *et al.^9^*. The 10-block contiguous and 10-block noncontiguous recombination libraries were designed and built using SCHEMA recombination^9^. Software packages for calculating SCHEMA energies are openly available at cheme.che.caltech.edu/groups/fha/Software.htm. Selected ChR variant genes were inserted into a constant vector backbone [pFCK from Addgene plasmid #51693^22^] with a CMV promoter, Golgi export trafficking signal (TS) sequence (KSRITSEGEYIPLDQIDINV)^5^, and fluorescent protein (mKate). All ChR variants contain the SpyTag sequence following the N-terminal signal peptide for the SpyTag/SpyCatcher labeling assays used to characterize ChR membrane localization^9,32^. The C1C2 parent for the recombination libraries is mammalian codon-optimized. For characterization in neurons, selected ChR variants [ChRger1, ChRger2, ChRger3, CoChR^11^, and hChR2(H134R)] were inserted into a pAAV-hSyn vector backbone [Addgene plasmid #26973], a pAAV-CamKIIa vector backbone [Addgene plasmid #51087], and a pAAV-CAG-DIO vector backbone [Addgene plasmid #104052]. In all backbones, each ChR was inserted with a Golgi export trafficking signal (TS) sequence (KSRITSEGEYIPLDQIDINV)^5^, and fluorescent protein (eYFP). ChR variant sequences used in this study are documented in **Dataset 2**. All selected ChR genes were synthesized and cloned in the pFCK mammalian expression vector by Twist Bioscience. HEK293T cells were transfected with purified ChR variant DNA using FuGENE^®^6 reagent according to the manufacturer’s (Promega) recommendations. Cells were given 48 hours to express the ChRs before photocurrent measurements. Imaging of ChR variants expression in HEK cells was performed using an Andor Neo 5.5 sCMOS camera and Micro-Manager Open Source Microscopy Software. Imaging of ChR expression in neuronal cultures and in brain slices was performed using a Zeiss LSM 880 confocal microscope and Zen software.

#### Primary neuronal cultures

Primary hippocampal neuronal cultures were prepped from C57BL/6N mouse embryos 16-18 days post-fertilization (E16-E18 Charles-River Labs) and cultured at 37 °C in the presence of 5% CO_2_ in Neurobasal media supplemented with glutamine and B27. Cells were transduced 3 − 4 days after plating with rAAV-PHP.eB packaging ChR2(H134R), ChRger1, ChRger2, or ChRger3. Whole-cell recordings were performed 10 − 14 days after transduction.

#### Patch-clamp electrophysiology

Whole-cell patch-clamp and cell-attached recordings were performed in transfected HEK cells, transduced neurons, and acute brain slices to measure light-activated inward currents or neuronal firing. For electrophysiological recordings, cultured cells were continuously perfused with extracellular solution at room temperature (in mM: 140 NaCl, 5 KCl, 10 HEPES, 2 MgCl_2_, 2 CaCl_2_, 10 glucose; pH 7.35) while mounted on the microscope stage. For slice recordings, 32 °C artificial cerebrospinal fluid (ACSF) was continuously perfused over slices. ACSF contained 127 mM NaCl, 2.5 mM KCl, 25 mM NaHCO_3_, 1.25 mM NaH_2_PO_4_, 12 mM *d*-glucose, 0.4 mM sodium ascorbate, 2 mM CaCl_2_, and 1 mM MgCl_2_ and was bubbled continuously with 95% oxygen / 5% CO_2_.

Patch pipettes were fabricated from borosilicate capillary glass tubing (1B150-4; World Precision Instruments) using a model P-2000 laser puller (Sutter Instruments) to resistances of 3–6 MΩ. Pipettes were filled with K-gluconate intracellular solution containing the following (in mM): 134 K gluconate, 5 EGTA, 10 HEPES, 2 MgCl_2_, 0.5 CaCl_2_, 3 ATP, and 0.2 GTP. Whole-cell patch-clamp and cell-attached recordings were made using a Multiclamp 700B amplifier (Molecular Devices), a Digidata 1440 digitizer (Molecular Devices), and a PC running pClamp (version 10.4) software (Molecular Devices) to generate current injection waveforms and to record voltage and current traces. Access resistance (*R*_a_) and membrane resistance (*R*_m_) were monitored throughout recording.

Patch-clamp recordings were done with short light pulses to measure photocurrents. Light pulse duration, wavelength, and power were varied depending on the experiment (as described in the text). Light pulses were generated using a Lumencor SPECTRAX light engine and quad band 387/485/559/649 nm excitation filter (SEMROCK, Part Number: FF01-387/485/559/649-25). To evaluate normalized green photocurrent, we measured photocurrent strength at three wavelengths: (red) 650 ± 13 nm LED with 643 − 656 nm filter, (green) 560 ± 25 nm LED with 547 − 572 nm filter, and (cyan) 485 ± 20 nm LED with 475 − 495 nm filter with a 0.5 s light pulse. Light intensity was matched for these measurements, with 485 nm light at 2.3 mW mm^-2^, 560 nm light at 2.8 mW mm^-2^, and 650 nm light at 2.2 mW mm^-2^. For full spectra measurements depicted in **Figure 3e**, we measured photocurrents at seven different wavelengths: (red) 650 ± 13 nm LED, (yellow) 586 ± 20 nm LED, (green) 560 ± 25 nm LED, (teal) 513 ± 22 nm LED, (cyan) 485 ± 20 nm LED, (blue) 438 ± 29 nm LED, and (violet) 395 ± 25 nm LED with a 0.5 s light pulse for each color. Light intensity is matched across wavelengths at 1.3 mW mm^-2^.

Photocurrents were recorded from cells in voltage clamp held at −60 mV. Neuronal firing was measured in current clamp mode with current injection for a −60 mV holding potential. For cell culture experiments, the experimenter was blinded to the identity of the ChR being patched but not to the fluorescence level of the cells. For acute slice recordings, the experimenter was not blinded to the identity of the ChR.

Electrophysiology data were analyzed using custom data-processing scripts written using open-source packages in the Python programming language to perform baseline adjustments, find the peak and steady state inward currents, perform monoexponential fits of photocurrent decay for off-kinetic properties, and quantify spike fidelity. Only cells with an uncompensated series resistance between 5 and 30 MΩ, *R*_m_ > 90 MΩ, and holding current >-150 pA (holding at −60 mV) were included in data analysis. The photocurrent amplitude was not adjusted for expression level since both expression and conductance contribute to the *in vivo* utility of the tool. However, comparisons of expression with photocurrent strength for all ChR variants tested are included in **Supplemental Figures 5-7**.

Plotting and statistical analysis were done in Python and GraphPad Prism 7.01. For statistical comparisons, we performed non-parametric Kruskal-Wallis test with Dunn’s multiple comparisons *post hoc* test.

#### AAV production and purification

Production of recombinant AAV-PHP.eB packaging pAAV-hSyn-X-TS-eYFP-WPRE, pAAV-CAG-DIO[X-TS-eYFP]-WPRE, and pAAV-CaMKIIa-X-TS-eYFP-WPRE (X = ChR2(H134R), ChRger1, ChRger2, and ChRger3) was done following the methods described in Deverman *et al*.^33^ and Challis *et al.^28^*. Briefly, triple transfection of HEK293T cells (ATCC) was performed using polyethylenimine (PEI). Viral particles were harvested from the media and cells. Virus was then purified over iodixanol (Optiprep, Sigma; D1556) step gradients (15%, 25%, 40% and 60%). Viruses were concentrated and formulated in phosphate buffered saline (PBS). Virus titers were determined by measuring the number of DNase I–resistant viral genomes using qPCR with linearized genome plasmid as a standard.

#### Animals

All procedures were approved by the California Institute of Technology Institutional Animal Care and Use Committee (IACUC). *Dat*-Cre mice (006660) and C57Bl/6J mice (000664) were purchased from Jackson Laboratory.

#### Intravenous injections, stereotactic injections, and cannula implantation

Intravenous administration of rAAV vectors was performed by injecting the virus into the retro-orbital sinus at viral titers indicated in the text. There were no observed health issues with animals after systemic injection of virus at the titers presented in the paper. Mice remain healthy >6 months after systemic delivery of ChR2 and ChRgers. With slice electrophysiology, we did not observe any indication of poor cell health due to viral-mediated expression, which we quantified by measuring the membrane resistance [*R*_m_], leak current [holding at −60 mV], and resting membrane potential. Local expression in the prefrontal cortex (PFC) was performed by direct stereotactic injection of 1 μl of purified AAV vectors at 5×10^12^ vg ml^-1^ targeting the following coordinates: anterior-posterior (AP), −1.7; media-lateral (ML), +/-0.5; and dorsal-ventral (DV), −2.2. For stimulation of the VTA, 300 μm outer diameter mono fiber-optic cannulae (Doric Lenses, MFC_300/330-0.37_6mm_ZF1.25_FLT) were stereotaxically implanted 200 μm above the VTA bilaterally targeted to the following coordinates: AP, −3.44 mm; ML, +/-0.48 mm; DV, 4.4 mm. For stimulation of the right secondary motor cortex (M2), 3 mm long, 400 μm mono fiber-optic cannulae (Doric Lenses, MFC_400/430-0.48_3mm_ZF1.25_FLT) were surgically secured to the surface of the skull above M2 (unilaterally) targeted to the following coordinates: AP, 1 mm; ML, 0.5 mm. The skull was thinned ~40 − 50% with a standard drill to create a level surface for the fiber-skull interface. Light was delivered from either a 447 nm or 671 nm laser (Changchun New Industries [CNI] Model with PSU-H-LED) via mono fiber-optic patch cable(s) (Doric Lenses, MFP_400/430/1100-0.48_2m_FC-ZF1.25) coupled to the fiber-optic cannula(e). Fiber-optic cannulae were secured to the skull with Metabond (Parkel, SKU S396) and dental cement.

Analysis of behavioral experiments was performed using the open-source MATLAB program OptiMouse^34^ to track mouse nose, body, and tail position while the mouse was running on the treadmill. Optogenetic intracranial self-stimulation was performed using a mouse modular test chamber (Lafayette Instruments, Model 80015NS) outfitted with an IR nose port (Model 80116TM).

#### Gaussian process modeling

Both the GP regression and classification modeling methods applied in this paper are based on work detailed in ref 8 and 23. For modeling, all sequences were aligned using MUltiple Sequence Comparison by Log-Expectation (MUSCLE) (https://www.ebi.ac.uk/Tools/msa/muscle/). For modeling, aligned sequences were truncated to match the length of the C1C2 sequence, eliminating N- and C-terminal fragments with poor alignment quality due to high sequence diversity (**Dataset 1** and **Dataset 2**). Structural encodings use the C1C2 crystal structure (3UG9.pdb) and assume that ChR chimeras share the contact architecture observed in the C1C2 crystal structure. For a given ChR, the contact map is simply a list of contacting amino acids with their positions. For example, a contact between alanine at position 134 and methionine at position 1 of the amino acid sequence would be encoded by [`(‘A134’), (‘M1’)`]. Both sequence and structural information were one-hot encoded. Regression models for ChR properties were trained to predict the logarithm of the measured properties. All training data was normalized to have mean zero and standard deviation one.

Gaussian process regression and classification models require kernel functions that measure the similarity between protein sequences. Learning involves optimizing the form of the kernel and its hyperparameters (**Supplemental Table 2**). The Matérn kernel was found to be optimal for all ChR properties (**Table 1**).

##### GP regression

In regression, the goal is to infer the value of an unknown function *f*(*x*) at a novel point *x*_*_ given observations *y* at inputs *X*. Assuming that the observations are subject to independent and identically distributed Gaussian noise with variance 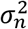, the posterior distribution of *f*_*_ = *f*(*x*_*_) for Gaussian process regression is Gaussian with mean

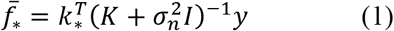

and variance

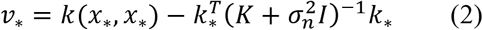

Where *K* is the symmetric, square covariance matrix for the training set: *K_ij_* = *k*(*x_i_*, *x_j_*) for *x_i_* and *x_j_* in the training set. *k*_*_ is the vector of covariances between the novel input and each input in the training set, and *k*_\**i*_ = *k*(*x*_*_, *x_i_*). The hyperparameters in the kernel functions and the noise hyperparameter *σ_n_* were determined by maximizing the log marginal likelihood:

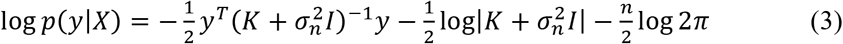

where *n* is the dimensionality of the inputs. Regression was implemented using open-source packages in the SciPy ecosystem^35–37^.

##### GP classification

In binary classification, instead of continuous outputs *y*, the outputs are class labels *y_i_* ∈ {+1, −1}, and the goal is to use the training data to make probabilistic predictions *π*(*x*_*_) = *p*(*y*_*_ = +1|*x*_*_). We use Laplace’s method to approximate the posterior distribution. Hyperparameters in the kernels are found by maximizing the marginal likelihood. Classification was implemented using open-source packages in the SciPy ecosystem^35–37^.

##### GP kernels for modeling proteins

Gaussian process regression and classification models require kernel functions that measure the similarity between protein sequences. A protein sequence *s* of length *L* is defined by the amino acid present at each location. This can be encoded as a binary feature vector *x_se_* that indicates the presence or absence of each amino acid at each position resulting in a vector of length 20*L* (for 20 possible amino acids). Likewise, the protein’s structure can be represented as a residue-residue contact map. The contact map can be encoded as a binary feature vector *x_st_* that indicates the presence or absence of each possible contacting pair. We used both the sequence and structure feature vectors by concatenating them to form a sequence-structure feature vector.

We considered three types of kernel functions *k*(*s_i_*, *s_j_*): polynomial kernels, squared exponential kernels, and Matérn kernels. These different forms represent possible functions for the protein’s fitness landscape. The polynomial kernel is defined as:

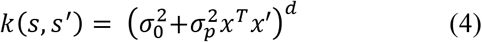

where *σ*_0_ and *σ_p_* are hyperparameters. We considered polynomial kernels with *d* = 3. The squared exponential kernel is defined as:

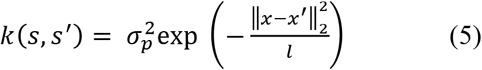

where *l* and *σ_p_* are also hyperparameters and | · |_2_ is the L2 norm. Finally, the Matérn kernel with 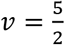 is defined as:

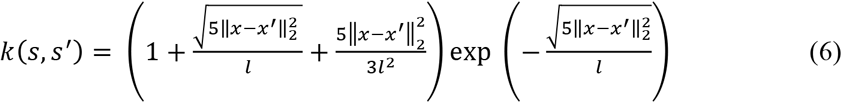

Where *l* is once again a hyperparameter.

##### L1 regression feature identification and weighting

We used L1 regression to identify residues and contacts in the ChR structure most important for each ChR functional property of interest. Using the concatenated sequence and structure binary feature vector for each of the training set ChR variants, we identified residues and contacts that covary. Each set of covarying residues and contacts was combined into a single feature. L1 linear regression was used to select the features that contribute most to each ChR functional property of interest. The level of regularization was chosen by maximizing the log marginal likelihood of the Gaussian process regression model trained on the features selected at that level of regularization. We then performed Bayesian ridge regression on the selected features using the default settings in scikit-learn^38^. Residues and contacts with the largest absolute Bayesian ridge linear regression weights were plotted onto the C1C2 structure (**Supplemental Figure 3 − 4**).

## Acknowledgements

We thank Twist Bioscience for synthesizing and cloning ChR sequences. We thank the Gradinaru and Arnold labs for helpful discussions. We also thank Dr. John Bedbrook for critical reading of the manuscript. This work was funded by the Beckman Institute for CLARITY, Optogenetics and Vector Engineering Research for technology development and broad dissemination: clover.caltech.edu (V.G.). This work was also supported by the National Institutes of Health (NIH) through NIH BRAIN grant R01MH117069 (V.G.) and SPARC grant OT2OD023848 (V.G), and by the Institute for Collaborative Biotechnologies through grant W911NF-09-0001 from the U.S. Army Research Office (F.H.A). The content of the information does not necessarily reflect the position or the policy of the Government, and no official endorsement should be inferred. C.N.B. is funded by Ruth L. Kirschstein National Research Service Awards F31MH102913. J.E.R. is supported by the Children’s Tumor Foundation (Young Investigator Award 2016-01-006).

## Author Contributions

C.N.B., K.K.Y., V.G., and F.H.A. conceptualized the project. C.N.B. coordinated all experiments and data analysis. C.N.B. and K.K.Y. built machine-learning models. C.N.B. performed construct design, cloning, and AAV production. C.N.B and J.E.R. conducted electrophysiology. C.N.B. and J.E.R. performed injections. J.E.R. performed fiber cannula implants and behavioral experiments. C.N.B. performed all data analysis. C.N.B. wrote the manuscript with input and editing from all authors. V.G. supervised optogenetics/electrophysiology parts of the project. F.H.A. supervised the protein engineering part of the project.

## Data availability

The authors declare that data supporting the findings of this study are available within the paper and its supplementary information files. Source data for classification model training are provided in **Dataset 1** and **Dataset 2**. Source data for regression model training are provided in **Dataset 2**.

## Code availability

Code used to train classification and regression models can be found at: https://github.com/fhalab/channels.

## Datasets

### Dataset 1

ChR sequence and photocurrent data from published sources including 19 natural ChR variants, 14 point-mutant ChR variants, and 28 recombination variants from various recombination libraries. The source of the photocurrent data is included (‘Reference’). When possible, we use references with side-by-side measurements of multiple ChRs. For modeling, all sequences were aligned and truncated to match the length of the C1C2 sequence (**Online methods**). The truncated and aligned sequences are included (‘Aligned_amino_acid_sequence’) as well as the full-length sequence (‘Amino_acid_sequence’).

### Dataset 2

ChR chimera sequences and functional properties for designed variants from our ChR recombination libraries^8,9^. Functional properties were tested in HEK cells. Measurements of peak and steady-state photocurrent (nA) with 485 nm light at 2.3 mW mm^-2^ (‘cyan_peak’ & ‘cyan_ss’), 560 nm light at 2.8 mW mm^-2^ (‘green_peak’ & ‘green_ss’), and 650 nm light at 2.2 mW mm^-2^ (‘red_peak’ & ‘red_ss’) are included. The maximum peak (‘max_peak’) and maximum steady-state (‘max_ss’) photocurrent (nA) obtained with any wavelength are included. Measurement of the time (ms) to reach 50% of the light-exposed photocurrent after light removal is included (‘kinetics_off’). The ratio of peak photocurrent with 560 nm light to maximum photocurrent was calculated per each cell and average for each ChR variant (‘norm_green’). Off-kinetics (‘kinetics_off’) and spectral properties (‘norm_green’) were only included for ChR variants with steady-state photocurrent strength >0.02 nA. Each ChR recombination variant has a chimera identity (‘block_ID’) beginning with either ‘c’ or ‘n’ to indicate the contiguous or non-contiguous library^8,9^ followed by 10 digits indicating the parent that contributes each of the 10 blocks (‘0’: CheRiff, ‘1’:C1C2, and ‘2’:CsChrimR). Each ChR variant’s number of mutations away from the nearest parent (‘m’) is included. For modeling, all sequences were aligned and truncated to match the length of the C1C2 sequence (**Online methods**). The truncated and aligned sequences are included (‘Aligned_amino_acid_sequence’) as well as the full-length sequence (‘Amino_acid_sequence’).

### Dataset 3

ChR variants predicted to localize and function. 1,161 ChR variants from the recombination libraries are above the 0.4 threshold for the product (‘pp’) of the predicted probabilities of localization (‘p_loc’) and function (‘p_func’). For all remaining variants (i.e., variants that we have not yet measured), we include the regression models’ prediction of peak photocurrent in nA (‘mu_peak_nA’), off-kinetics (time [ms] to reach 50% of the light-exposed photocurrent after light removal; ‘mu_kin_ms’), and normalized photocurrent with 560 nm light (‘mu_green’). We also include ChR variants’ amino acid and nucleic acid sequences.

### Dataset 4

Limited set of amino acid residues and structural contacts important for model predictions identified with L1-regularized linear regression. The relative importance (‘weight’) of these sequence and structural features is learned using Bayesian ridge regression. We found a different limited set of features for each of the three functional properties of interest (‘norm_green’, ‘off_kinetics’, and ‘peak_photocurrent’). Features are either amino acid residues (i.e. a sequence feature [‘seq’]) or contacts. The feature position is indicated with numbering according to the aligned and truncated ChR sequence. We also include the parental features at each position with numbering according the parental sequence. Highly-weighted features highlighted in color in **Supplemental Figure 3-4** are indicated by their corresponding color. Features not highlighted in **Supplemental Figure 3-4** are listed as gray.

