## Supplemental_Material for "Machine learning-guided channelrhodopsin engineering enables minimally-invasive optogenetics"

### Supplemental Figure 1

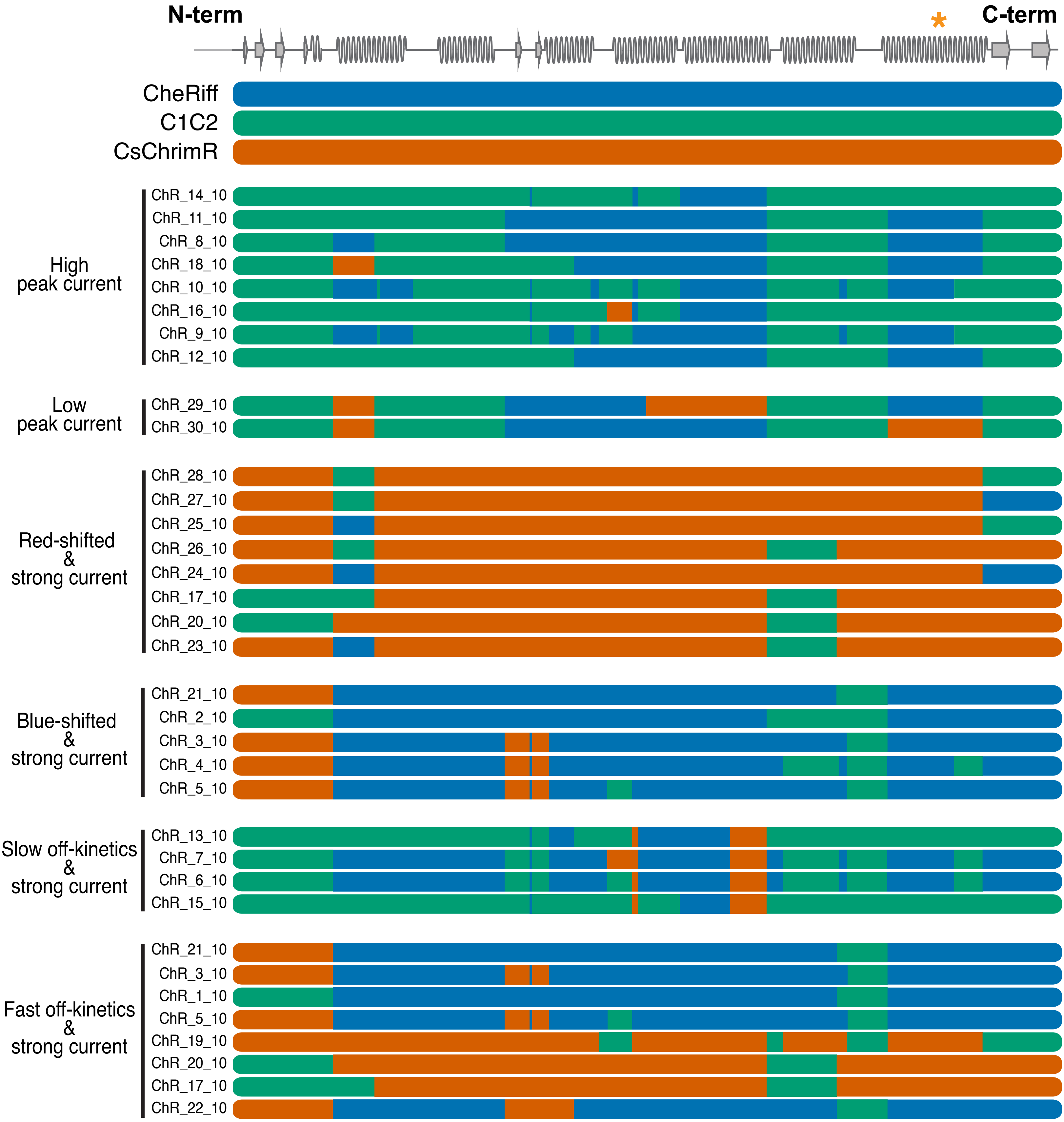

**Supplemental Figure 1**. Thirty model-predicted ChR chimeras aligned with the three parents and the secondary structure. Blocks of ChR chimeras are colored according to which parent each block came from. CsChrimR is red, CheRiff is blue, and C1C2 is green. (*) highlights the Schiff base. ChRs are divided into categories based on their predicted properties. Twenty-eight ChR chimeras are predicted to be optimized for one or more properties. Two ChR chimeras are predicted to be non-optimal and produce low currents. A number of chimeras appear twice because they were optimal for multiple categories.

### Supplemental Figure 2

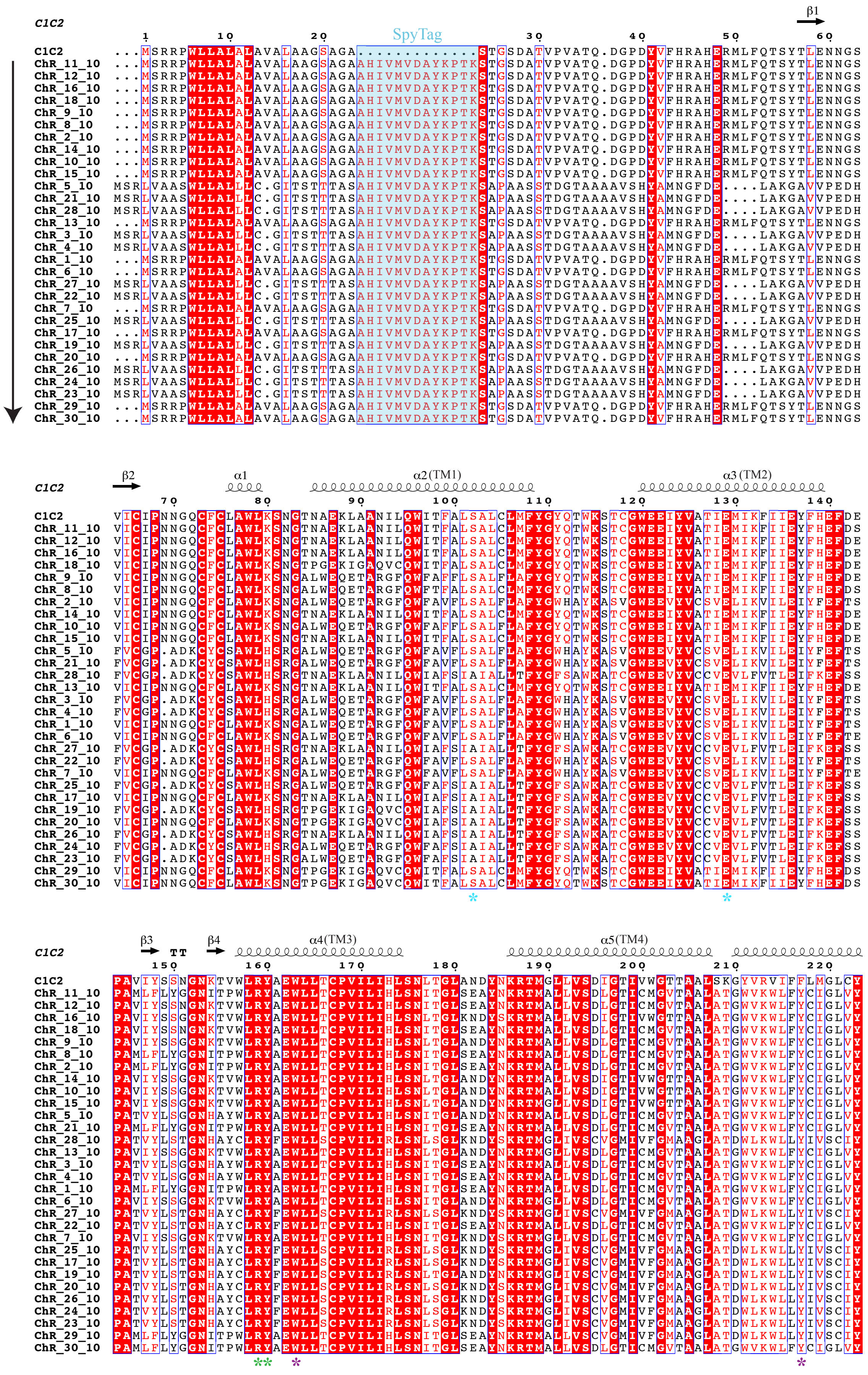

### Supplemental Figure 2

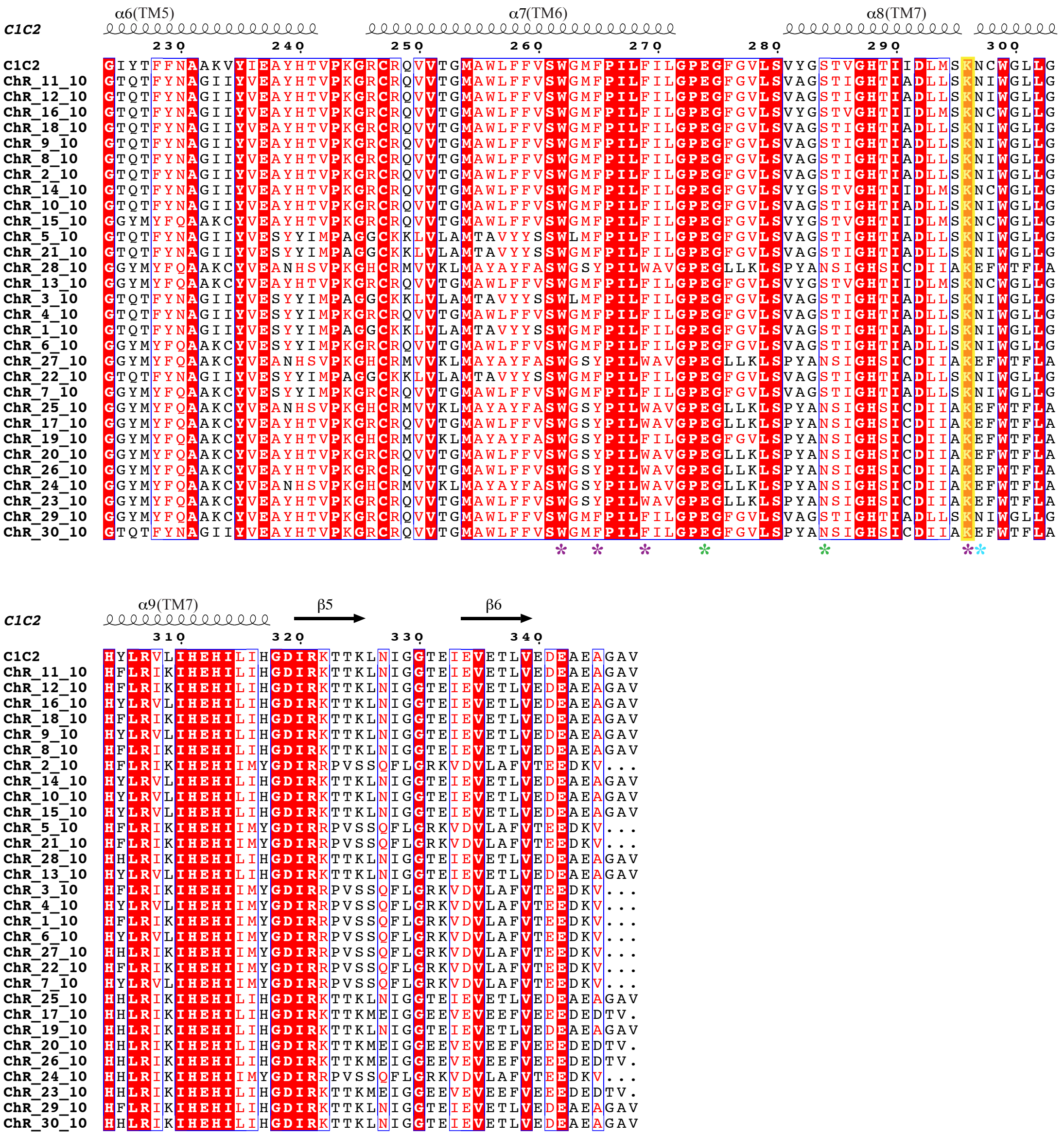

**Supplemental Figure 2**. The sequence alignment between C1C2 and designer ChRs was created using ClustalΩ and visualized using ENDscript[1]. Designer ChRs are arranged under the C1C2 sequence in order of decreasing photocurrent strength (ChR_11_10 has the strongest photocurrents while ChR_30_10 has the weakest photocurrents). Secondary structure elements for C1C2 are shown as coils (α: α-helices) and arrows (β-strands). "TT" represents turns. Identical and conservatively substituted residues are highlighted in red (outlined in blue box). Light-blue asterisks under the alignment indicate the three residues that form the internal gate. Purple and green asterisks under the alignment indicate the residues that form the conserved hydrophobic retinal-binding pocket and the conserved cluster at the extracellular vestibule of the cation-conducting pathway, respectively. The lysine residue involved in the Schiff base is highlighted in yellow shading. The SpyTag sequence is highlighted in light blue shading.

**Supplemental Figure 3**

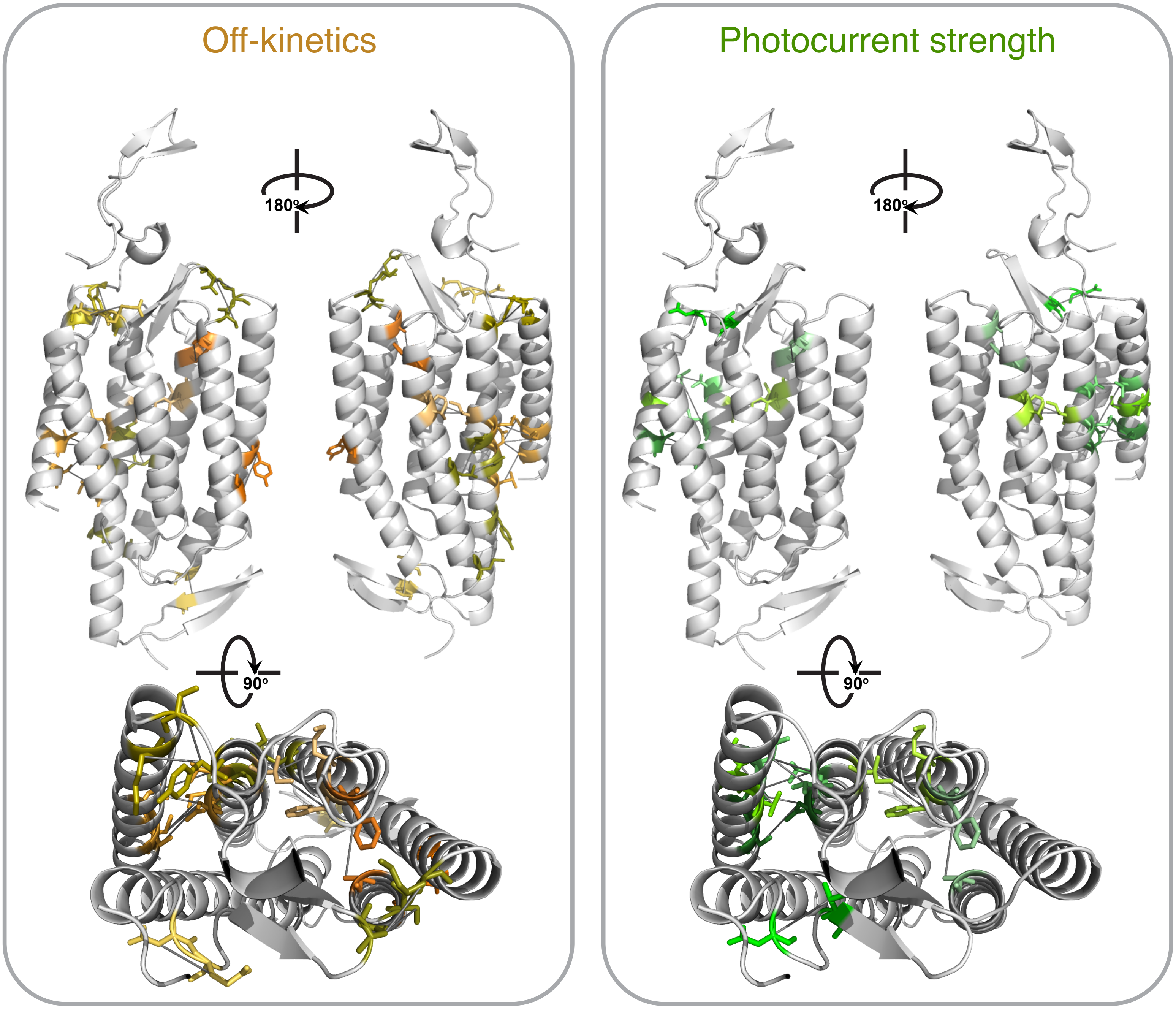

**Supplemental Figure 3**. Specific residues (amino-acid sticks) and contacts (dark gray lines) most important for model prediction of off-kinetics and photocurrent strength overlaid on the C1C2 crystal structure in light gray (3ug9.pdb). Specific residues and contacts are included in **Dataset 4**.

**Supplemental Figure 4**

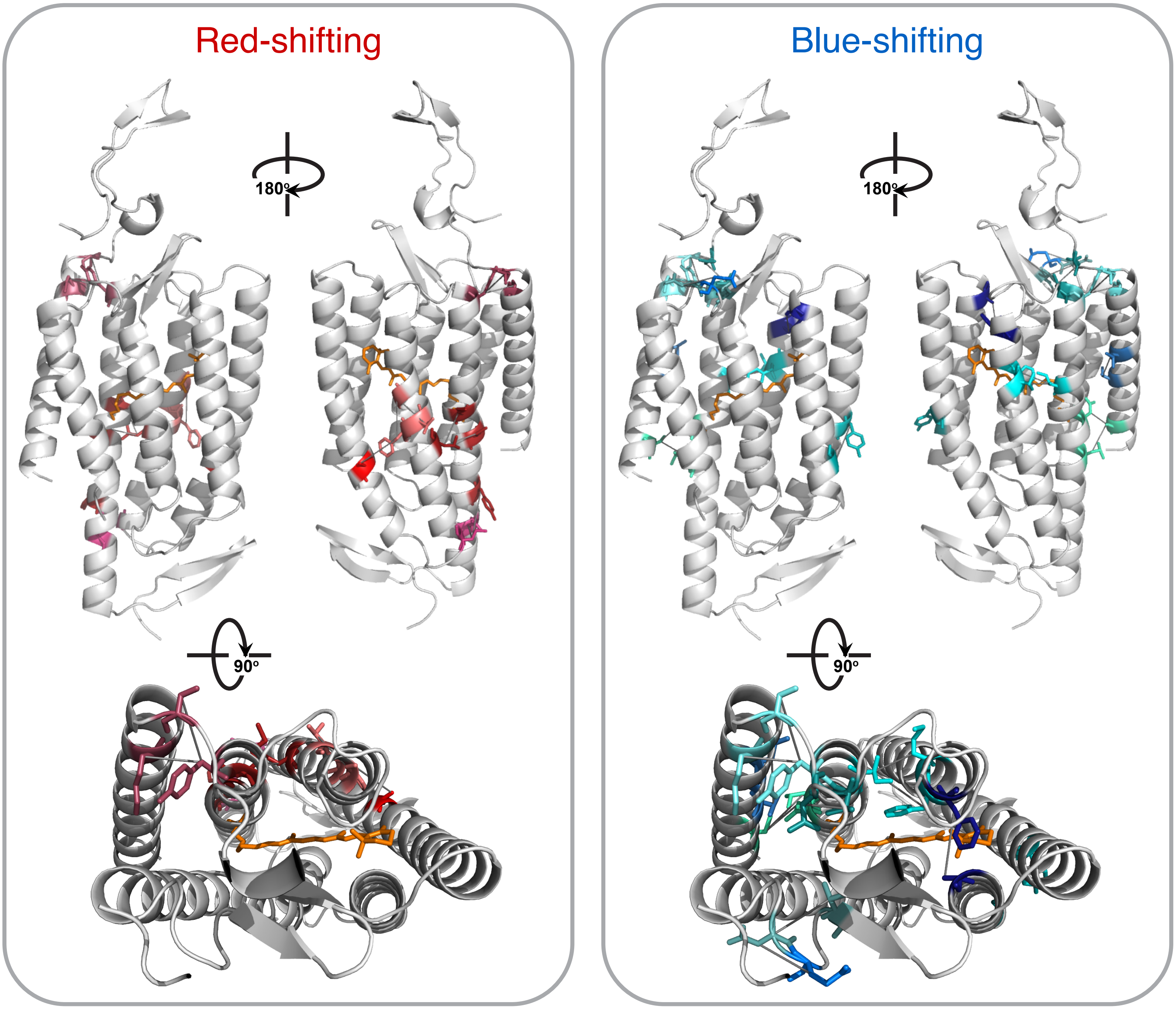

**Supplemental Figure 4**. Specific residues (amino-acid sticks) and contacts (dark gray lines) most important for model prediction of red-shifted light sensitivity and blue-shifted light sensitivity overlaid on the C1C2 crystal structure in light gray (3ug9.pdb). Specific residues and contacts are included in **Dataset 4**.

### Supplemental Figure 5

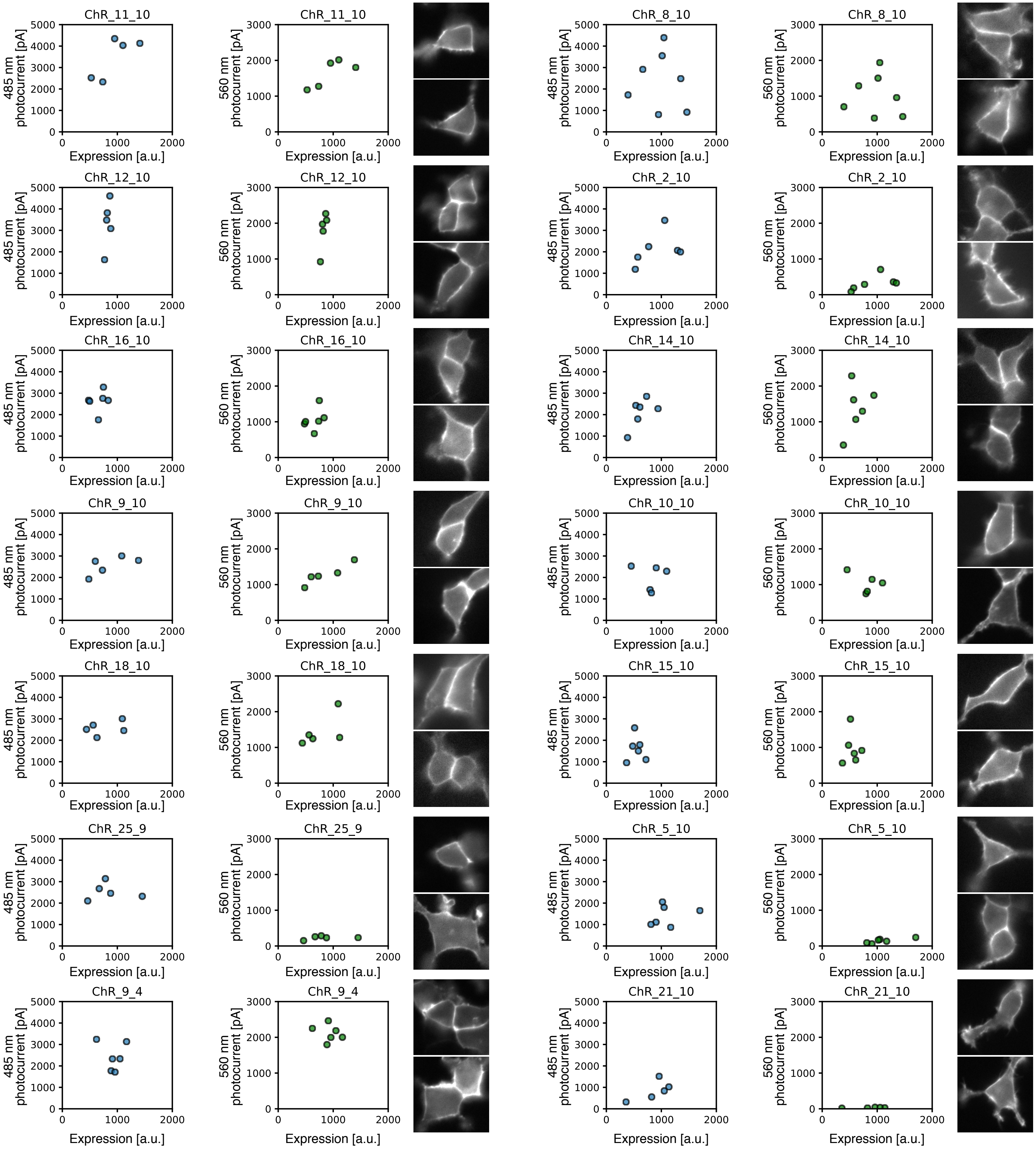

### Supplemental Figure 5

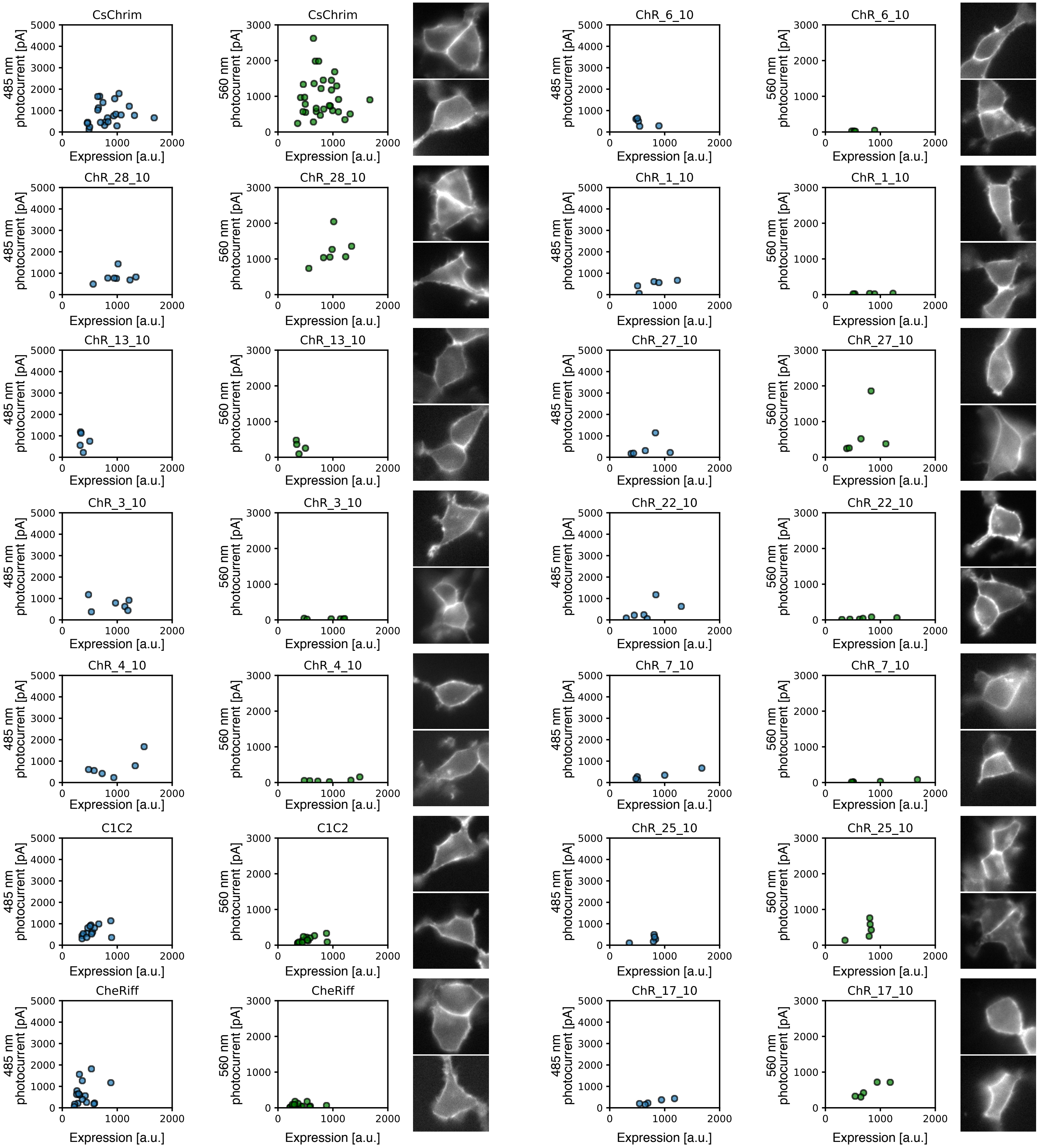

### Supplemental Figure 5

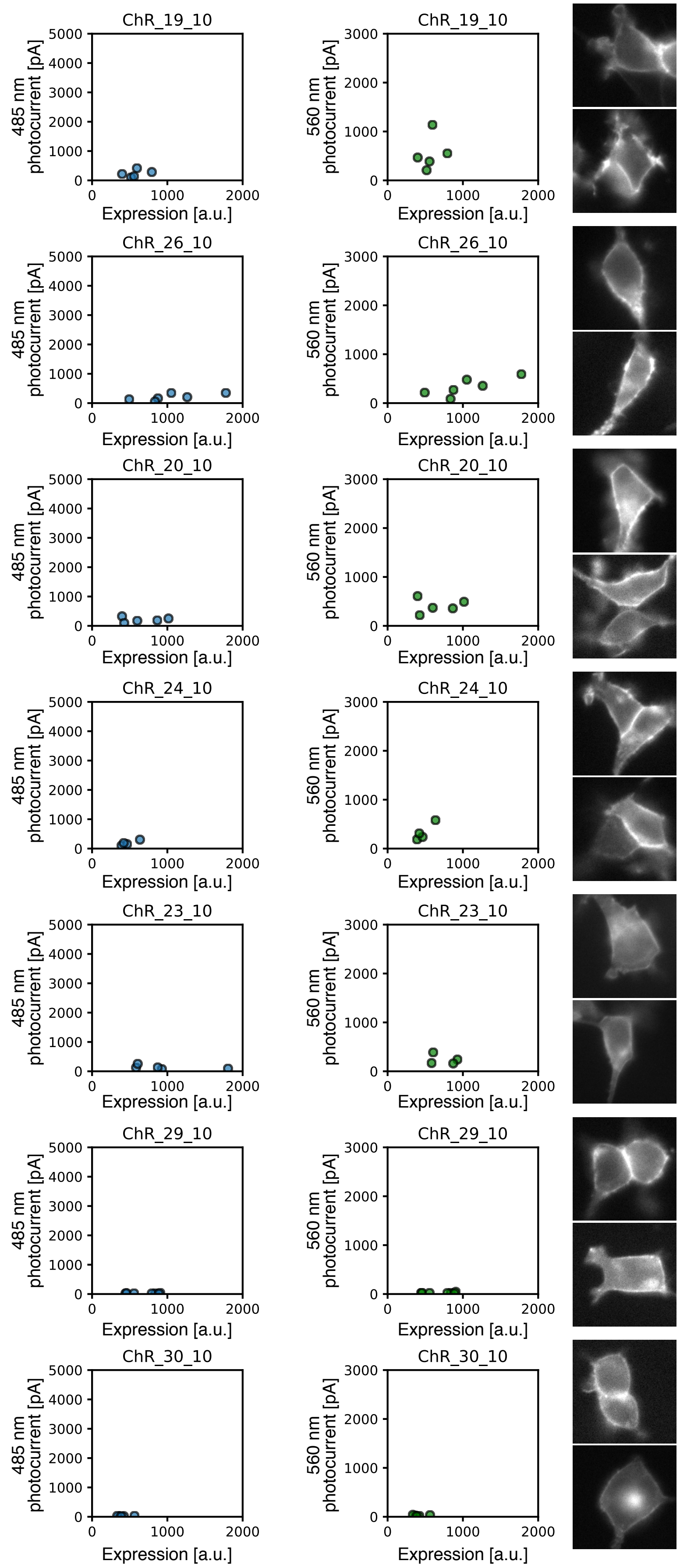

**Supplemental Figure 5**. Correlation between photocurrent strength and expression of ChR variants. Plot of measured photocurrent strength versus expression in HEK cells for each ChR variant. Each point is an individual cell. For each variant, images of two representative cells shows localization. To highlight ChR variant localization patterns, contrast in each image was adjusted so that localization can be compared for both high-expressing and low-expressing variants. Thus, images are not contrast matched and fluorescence brightness is not an indicator of relative expression level across variants.

#### Supplemental Figure 6

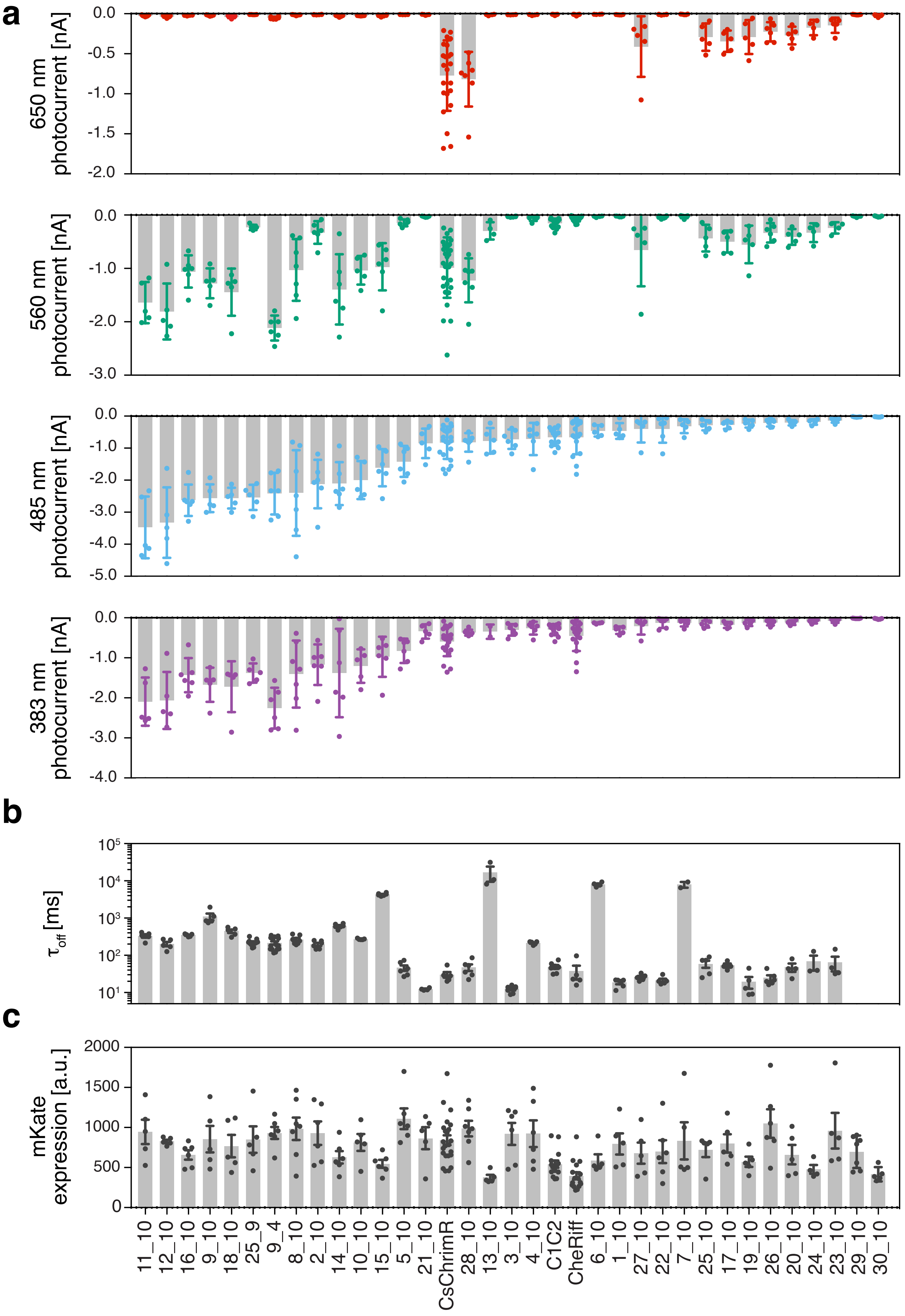

**Supplemental Figure 6**. Model-predicted ChRs exhibit a large range of functional properties often far exceeding the parents. This figure shows data depicted in **Figure 3** with ChR properties (photocurrent strength, off-kinetics, and expression level) aligned for each ChR variant for easy comparison. In all plots, each point is an individual cell. (**a**) Designer ChR measured peak photocurrent with different wavelengths of light in HEK cells (*n* = 4–8 cells, see **Dataset 2**). 383 nm light at 1.5 mW $mm^{-2}$, 485 nm light at 2.3 mW $mm^{-2}$, 560 nm light at 2.8 mW $mm^{-2}$, and 650 nm light at 2.2 mW $mm^{-2}$. (**b**) Designer ChR off-kinetics decay rate ($\tau_{off}$) following a 1 ms exposure to 485 nm light at 2.3 mW $mm^{-2}$ (*n* = 4–8 cells, see **Dataset 2**). (**c**) ChR-mKate measured expression level (*n* = 4–8 cells).

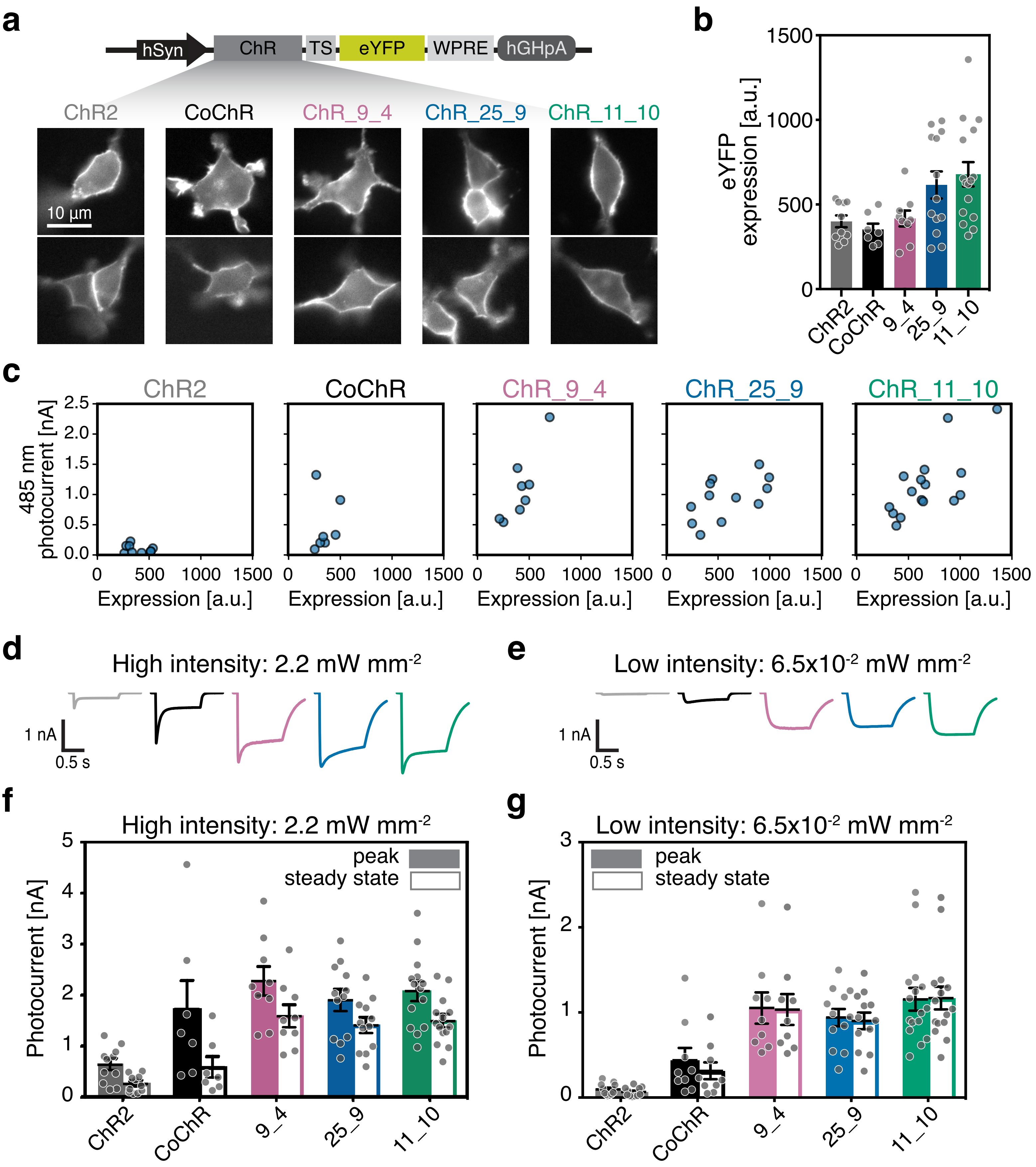

a
hSyn
ChR
TS
eYFP
WPRE
hGHpA
ChR2
CoChR
ChR_9_4
ChR_25_9
ChR_11_10
10 µm
b
eYFP expression [a.u.]
1500
1000
500
0
ChR2
CoChR
9_4
25_9
11_10
c
485 nm photocurrent [nA]
Expression [a.u.]
d
High intensity: 2.2 mW mm-2
1 nA
0.5 s
e
Low intensity: 6.5x10-2 mW mm-2
f
High intensity: 2.2 mW mm-2
Photocurrent [nA]
peak
steady state
g
Low intensity: 6.5x10-2 mW mm-2

**Supplemental Figure 7**. (**a**) Construct design for each ChR tested with a TS sequence, eYFP, and WPRE under the hSyn promoter. Representative cells show expression and localization of each ChR variant. (**b**) Measured expression of ChR-eYFP for each ChR construct. (**c**) Plot of measured photocurrent strength versus expression in HEK cells for each ChR variant. Each point is an individual cell. Current trace after 1 s light exposure for ChR variants with both (**d**) high-intensity and (**e**) low-intensity light. Peak and steady-state photocurrent comparison between high-performance ChRs, ChR2(H134R), and CoChR with both (**f**) high-intensity and (**g**) low-intensity light. Multiple HEK cells were recorded from for each ChR: ChR2(H134R), $n = 11$ cells; CoChR, $n = 7$ cells; ChR_9_4, $n = 9$ cells; ChR_25_9, $n = 12$ cells; ChR_11_10, $n = 16$ cells. Plotted data are mean ± SEM and each point is an individual cell. There is a significant difference between the high-performance ChR variants and ChR2(H134R) with low intensity light ($P = 0.0002$ for ChR_9_4, $P = 0.0001$ for ChR_25_9, and $P < 0.0001$ for ChR_11_10); Kruskal-Wallis test with Dunn's *post hoc* test, with ChR2 as a reference. There is a significant difference between the high-performance ChR variants and CoChR with low intensity light ($P = 0.04$ for ChR_9_4, $P = 0.04$ for ChR_25_9, and $P = 0.004$ for ChR_11_10); Kruskal-Wallis test with Dunn's *post hoc* test, with CoChR as a reference. Top variants, ChR_9_4, ChR_25_9, and ChR_11_10 are named ChRger1, ChRger2, and ChRger3 in subsequent figures.

**Supplemental Figure 8**

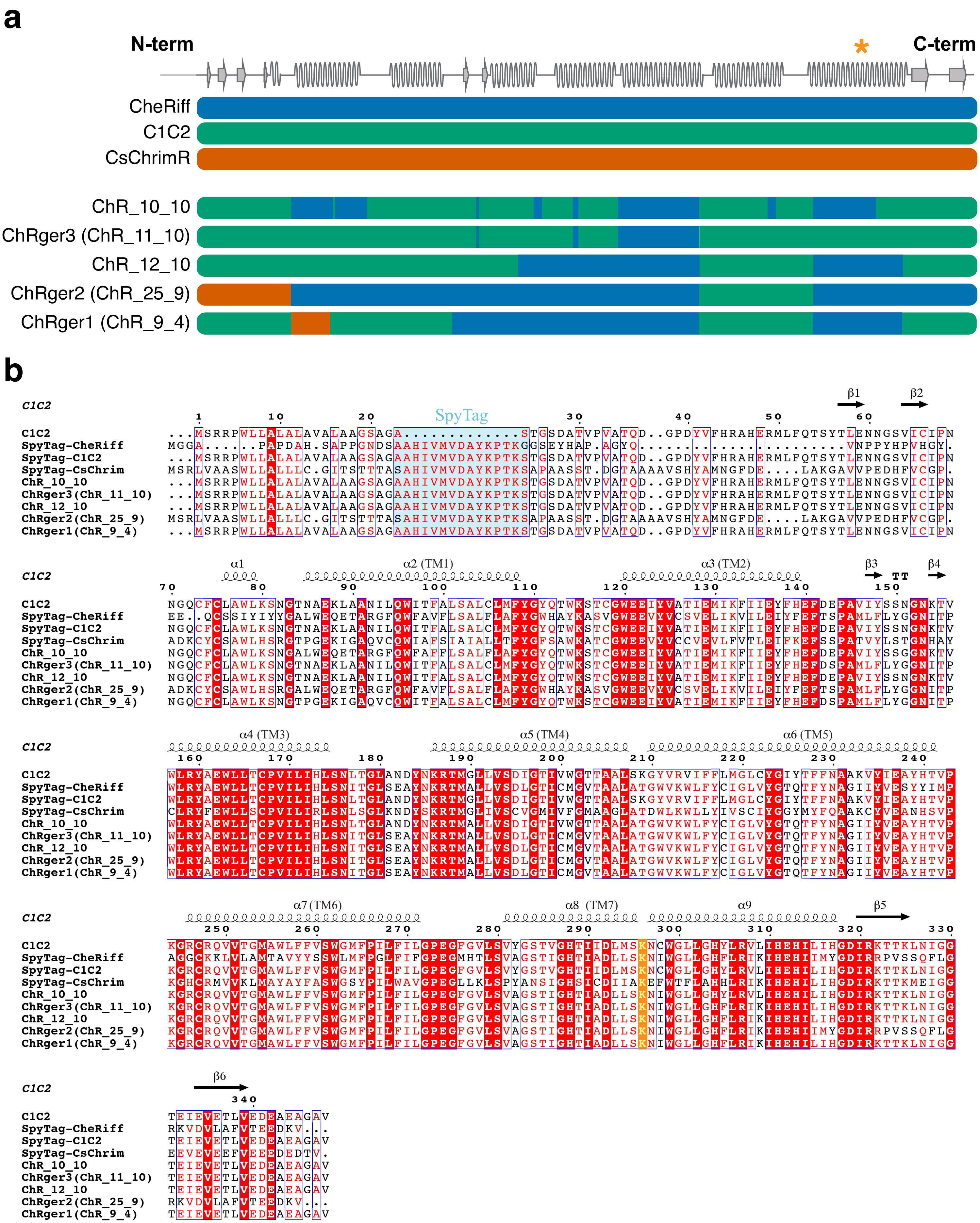

**Supplemental Figure 8**. (**a**) Top five designer ChRs predicted by the machine-learning models aligned with the three parents and the secondary structure. (*) highlights the Schiff base. Blocks of ChR chimeras are colored according to which parent each block came from. CsChrimR is red, CheRiff is blue, and C1C2 is green. (**b**) Sequence alignment between parents and top five designer ChRs was created using ClustalΩ and visualized using ENDscript[1]. Secondary structure elements for C1C2 are shown as coils (α: α-helices) and arrows (β-strands). "TT" represents turns. Identical and conservatively substituted residues are highlighted in red (outlined in blue box). The lysine residue involved in the Schiff base is highlighted in yellow shading. The SpyTag sequence is highlighted in light blue shading.

**Supplemental Figure 9**

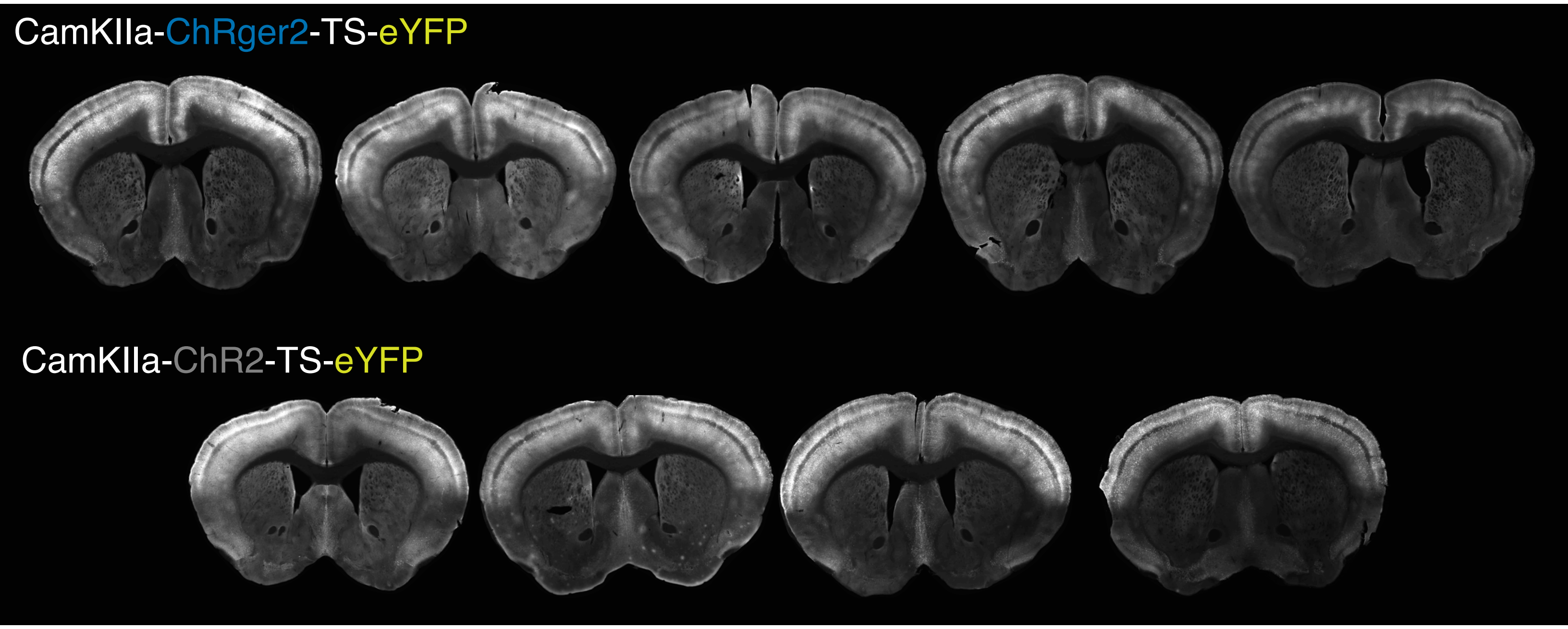

**Supplemental Figure 9**. Coronal slices show expression throughout cortex after systemic delivery of rAAV-PHP.eB packaged CaMKIIa ChRger2-TS-eYFP (top row; *n* = 5 animals) or ChR2(H134R)-TS-eYFP (bottom row; *n* = 4 animals) into wild type animals. These animals were used for behavioral experiments depicted in **Figure 5b**. Observable tissue damage occurred during tissue collection and processing. Virus titer, $5x10^{11}$ vg/mouse.

#### Supplemental tables

**Supplemental Table 1.** List of different constructs made for validation of the ChRgers.

| Vector | Insert (X) | Virus tested |
|---|---|---|
| **pAAV-hSyn-X-TS-eYFP-WPRE** | hChR2(H134R) | Yes |
| | ChRger1 | |
| | ChRger2 | |
| | ChRger3 | |
| **pAAV-CaMKIIa-X-TS-eYFP-WPRE** | hChR2(H134R) | Yes |
| | ChRger1 | |
| | ChRger2 | |
| | ChRger3 | |
| **pAAV-CAG-DIO[X-TS-eYFP]-WPRE** | hChR2(H134R) | Yes |
| | ChRger1 | |
| | ChRger2 | |
| | ChRger3 | |

**Supplemental Table 2.** GP regression model hyperparameters for each ChR property of interest for the Matérn kernel.

| Model type | ChR property | Noise hyperparameter: $\sigma_n^2$ | Length hyperparameter: $l$ |
|---|---|---|---|
| **GP regression** | current strength | 0.04848652 | 19.65389071 |
| **GP regression** | off-kinetics | 0.02902597 | 19.72715834 |
| **GP regression** | wavelength sensitivity | 0.10927067 | 37.7883682 |

#### Supplemental videos

**Supplementary Video 1.** ChRger2-expressing mouse running on a treadmill while receiving optogenetic stimulation exhibits clear left-turning behavior [10 Hz stimulation with 5 ms 447 nm light pulse (20 mW)]. Minimally-invasive, systemic delivery of rAAV-PHP.eB packaged CaMKIIa ChRger2-TS-eYFP ($5x10^{11}$ vg/mouse) into wild type (WT) animals coupled with surgically secured 2 mm long, 400 µm fiber-optic cannula guide to the surface of the skull above the right M2 that had been thinned to create a level surface for the fiber-skull interface (~40 – 50%). Video shows multiple runs with the same animal.

**Supplementary Video 2.** ChR2(H134R)-expressing mouse running on a treadmill while receiving optogenetic stimulation does not exhibit left-turning behavior [10 Hz stimulation with 5 ms 447 nm light pulse (20 mW)]. Minimally-invasive, systemic delivery of rAAV-PHP.eB packaged CaMKIIa ChR2(H134R)-TS-eYFP ($5x10^{11}$ vg/mouse) into wild type (WT) animals coupled with surgically secured 2 mm long, 400 µm fiber-optic cannula guide to the surface of the skull above the right M2 that had been thinned to create a level surface for the fiber-skull interface (~40 – 50%). Video shows multiple runs with the same animal.

1 Robert, X. & Gouet, P. Deciphering key features in protein structures with the new ENDscript server. *Nucleic acids research* **42**, W320-324, doi:10.1093/nar/gku316 (2014).
